# Cell wall synthesis and remodeling dynamics determine bacterial division site architecture and cell shape

**DOI:** 10.1101/2021.10.02.462887

**Authors:** Paula P. Navarro, Andrea Vettiger, Virly Y. Ananda, Paula Montero Llopis, Christoph Allolio, Thomas G. Bernhardt, Luke H. Chao

## Abstract

The bacterial division apparatus builds daughter cell poles by catalyzing the synthesis and remodeling of the septal peptidoglycan (sPG) cell wall. Understanding of this essential process has been limited by the lack of native three-dimensional visualization of developing septa. Here, we used state-of-the-art cryogenic electron tomography (cryo-ET) and fluorescence microscopy to understand the division site architecture and sPG biogenesis dynamics of the Gram-negative bacterium *Escherichia coli*. Our results with mutant cells altered in the regulation of sPG biogenesis revealed a striking and unexpected similarity between the architecture of *E. coli* septa with those from Gram-positive bacteria, suggesting a conserved morphogenic mechanism. Furthermore, we found that the cell elongation and division machineries are in competition and that their relative activities determine the shape of cell constrictions and the poles they form. Overall, our results highlight how the activity of the division system can be modulated to generate the diverse array of morphologies observed in the bacterial domain.

**Highlights:**

- The division site architecture of *E. coli* can be modulated to resemble that of diverse bacteria.
- Cell wall degradation at the division site activates septal cell wall synthesis.
- Assembly of the cytoskeletal ring at the division site is modulated by cell wall remodeling.
- Balance between the activities of the elongation and division systems modulates cell shape.

## Introduction

Bacterial cells are typically surrounded by a multi-layered cell envelope that varies in complexity depending on the species^1^. Organisms classified as Gram-positive (monoderm) possess a single membrane surrounded by a thick cell wall. Bacteria belonging to the Gram-negative (diderm) class have a thinner wall layer to which a second membrane called the outer membrane (OM) is attached. Their cell wall is therefore contained in an aqueous compartment called the periplasm that is sandwiched by the inner (cytoplasmic) membrane (IM) and the OM^2^. Understanding the mechanisms required for the biogenesis of bacterial envelopes during cell growth and division has been a long-standing and fundamental goal of microbiology. The process is also the target of many of our most widely used antibiotics. As such, studies of cell surface assembly are of great practical significance due to the enabling insights they provide for the development of new antibacterial treatments needed to combat the growing problem of drug-resistant infections.

The cell wall is the layer of the envelope that determines cell shape^3^. It is also critical for fortifying the cytoplasmic membrane against osmotic lysis. The wall is assembled from the heteropolymer peptidoglycan (PG), which consists of glycan chains with repeating disaccharide units of N-acetylglucosamine (GlcNAc) and N-acetylmuramic acid (MurNAc). Short peptides are attached to each MurNAc sugar and are used to form amide crosslinks between adjacent glycans, generating a covalently closed mesh around the cytoplasmic membrane. Because the PG layer completely envelops the cell, its expansion and remodeling are tightly coordinated with the cell cycle^3^.

Newborn rod-shaped cells like *Escherichia coli* first lengthen their cell body through the action of the elongation machinery (Rod complex, elongasome), which synthesizes and incorporates new PG material at dispersed locations throughout the cylinder^3^. Cell division is then initiated by the coalescence of treadmilling polymers of the tubulin-like FtsZ protein into a structure called the Z-ring at midcell^4,5^. The Z-ring is peripherally associated with the IM and its formation ultimately results in the recruitment of dozens of proteins to the prospective division site to assemble the envelope-spanning division machinery called the divisome^5,6^. A major function of this apparatus is to promote the localized synthesis of PG to generate the cross-wall/septum that divides the daughter cell compartments^3^. The septal PG (sPG) produced by the divisome initially connects the daughters such that it must be carefully processed to separate the newly formed cells and complete the division process^3^.

Our understanding of envelope biogenesis during cell division has been greatly influenced by electron micrographs of developing septa^7^. One of the most striking observations from these images is the extent to which the septal architectures differ between organisms^8^. Some of the best images of septa from a Gram-positive bacterium come from cryogenic electron microscopy of frozen vitrified sections (CEMOVIS) of *Staphylococcus aureus*^9,10^. In the septa of these cells, two tracks of PG material appear to be connected to the lateral cell wall, forming an electron dense π structure with the two tracks located between the invaginating membrane. Other Gram-positive bacteria like *Bacillus subtilis* are only observed to produce a single, thick layer of cell wall within their septa^11–14^. In contrast to the flat unconstricted side wall present at Gram-positive division sites, conventional electron microscopy (EM) of fixed and stained cells and whole-cell cryo-electron tomography (cryo-ET) imaging of *E. coli* and *Caulobacter crescentus* cells have visualized a V-shaped constriction in which all three envelope layers are invaginating together^15–17^. Although the general presence of sPG can be observed in some images, the existence of sample preparation artifacts and the low signal-to-noise ratio in conventional EM and cryo-ET images, respectively, have prevented a clear visualization of its structural arrangement. However, notable morphological differences between the division sites of *E. coli* and *C. crescentus* were evident in these images. Division sites from *C. crescentus* displayed very tight coordination between the invagination of the three envelope layers in all cells that were imaged^16^. They were never observed to form what would be considered a classical crosswall septum in which a band of sPG bisects the cytoplasm. By contrast, some images of *E. coli* showed a coordinately constricted envelope whereas others showed a partial crosswall where the IM and PG layers seem more invaginated than the OM^15^. Whether these different septal architectures reflect sample preparation artifacts, fundamental differences in the division mechanism between bacteria or arise from changes in the spatiotemporal regulation of conserved processes remains a major outstanding question.

To investigate the underlying mechanisms responsible for generating different division site morphologies, we imaged the *in situ* ultrastructure of the division site and the dynamics of cell envelope constriction in wild-type *E. coli* and several mutant derivatives with altered divisome function using cryo-ET and fluorescence microscopy. For the cryo-ET analysis, we needed to overcome the major sample thickness limitation of this imaging method in which samples > 300 nm in depth preclude the generation of distinguishable contrast features by EM^18,19^. This thickness limit is incompatible with the width of *E. coli* and most other well-studied bacterial species that have cell diameters near 1 μm^20^. The development of cryo-focused ion beam (cryo-FIB) milling^21–27^ made this imaging possible by allowing us to generate thin, artifact-free lamellae of frozen-hydrated *E. coli* cells in various stages of division that are within the optimal thickness regime for cryo-ET.

In this work, we used cryo-FIB milling and cryo-ET for the three-dimensional (3D) visualization of *E. coli* septa and the sPG layer with unprecedented detail, revealing several new ultrastructural features. Micrographs of wild-type cells showed that constriction of the IM and OM was well coordinated at the start of division, but that the distance between the membranes grew in cells at later stages of division due to much deeper invagination of the IM. Live-cell imaging with fluorescent markers for the IM and OM showed that this change in morphology arises from a more rapid constriction of the IM relative to the OM. Importantly, mutants impaired for sPG synthesis maintained coordination between IM and OM invagination throughout the division process and had division site architectures that resembled the shallowly constricted ultrastructure of *C. crescentus*. Moreover, mutants defective for sPG remodeling were shown to be largely blocked for OM constriction and their septa as well as those of wild-type cells were found to contain lateral bridge structures and separate parallel plates of sPG reminiscent of septa from *S. aureus*. In addition to tracking IM and OM dynamics, we also monitored the rates of sPG synthesis and remodeling at the division sites of the various mutants and correlated them with the rates of lateral wall biogenesis, division site ultrastructure, and the shape of daughter cell poles. Our results revealed that cell wall degradation at the division site plays a role in both the activation of sPG synthesis and Z-ring condensation, suggesting communication between the PG synthesis machinery on one side of the membrane and the cytoskeleton on the other. Furthermore, this analysis provided compelling evidence that the cell elongation and division machineries are in competition for shared PG precursors and that the relative activities of the two systems determines the shape of division site constrictions and the poles they form. Overall, our results highlight the plasticity of the division system and how modulation of its cell wall synthesis and remodeling activities can help generate the array of cell and polar morphologies observed in the bacterial domain.

## Results

### *In situ* architecture of the *E. coli* cell envelope at the division site

Most of the available images of the division site architecture in *E. coli* come from conventional EM micrographs of fixed and stained cells^28^. Although these images have given us a general idea of how the cell envelope is remodeled during division to build the daughter cell poles, the division site ultrastructure observed depended greatly on the fixation conditions used. Some treatments yielded division sites with a uniformly constricted architecture, whereas others captured cells with a partial crosswall septum composed of a fold of IM sandwiching a stretch of sPG from which the OM was excluded^15,29^. In contrast, whole-cell cryo-ET images of *E. coli* grown in nutrient poor conditions to reduce their size have thus far only shown uniformly constricted septa without a clear view of the sPG layer^17^. The variation in septal architecture based on the imaging method or fixation condition used has made it difficult to draw definitive conclusions about which architecture represents the native state of the division site. We therefore reinvestigated the ultrastructure of the *E. coli* division site using cryo-FIB milling to obtain thin (∼150-200 nm thick) lamellae of dividing cells for subsequent *in situ* cryo-ET imaging (**Fig. S1**). A total of 9 tilt-series of wild-type cells were acquired, aligned, and 3D reconstructed (**Table S1**). Additionally, to gain better visualization of sPG, non-linear anisotropic diffusion (NAD) filtering was applied to denoise the cryo-electron tomograms. Using the measured distance between the leading edge of the IM constriction for each image, they were grouped into three classes representing what appeared to be different stages of division: (i) constriction, (ii) septation, or (iii) cytokinesis (**Fig. 1A**). Cells with an IM-IM distance of around 500 nm were classified as being in the constriction phase. They had a V-shaped constriction with a relatively uniform invagination of the two membranes and an indented mesh of PG (**Fig. 1A-B, and S2A**). The second category of cells had a much smaller IM-IM distance (50 nm). They displayed a partial septum in which the IM was observed to be more deeply invaginated than the OM with an average difference of 134.1 (± 21) nm (mean ± SD) in their measured OM-OM versus IM-IM distance (**Fig. 1A-B, and S2B**). Strikingly, the denoised tomograms showed an elongated, triangular wedge of PG filling the gap between the two membranes (**Fig. 1B**). This structure is reminiscent of the lateral bridge in *S. aureus* septa connecting the plates of sPG^10^, except in this case a network of electron densities is visible connecting the two plates of PG that will become the daughter cell polar PG caps. In cells at the final stage of cytokinesis, IM fission was observed to be completed, leaving cells connected by the two interwoven sPG plates. The OM of these cells displayed deep constrictions with the leading edge of the membrane pointing between the two sPG plates, presumably poised to traverse the remaining OM-OM distance of 100-250 nm to complete division (**Fig. 1A-B, S2C and Video S1**). No OM blebs or cytoplasmic mesosome-like structures were observed at the division site in the cryo-ET data like those observed previously by conventional EM^15,28^.

**Figure 1:**
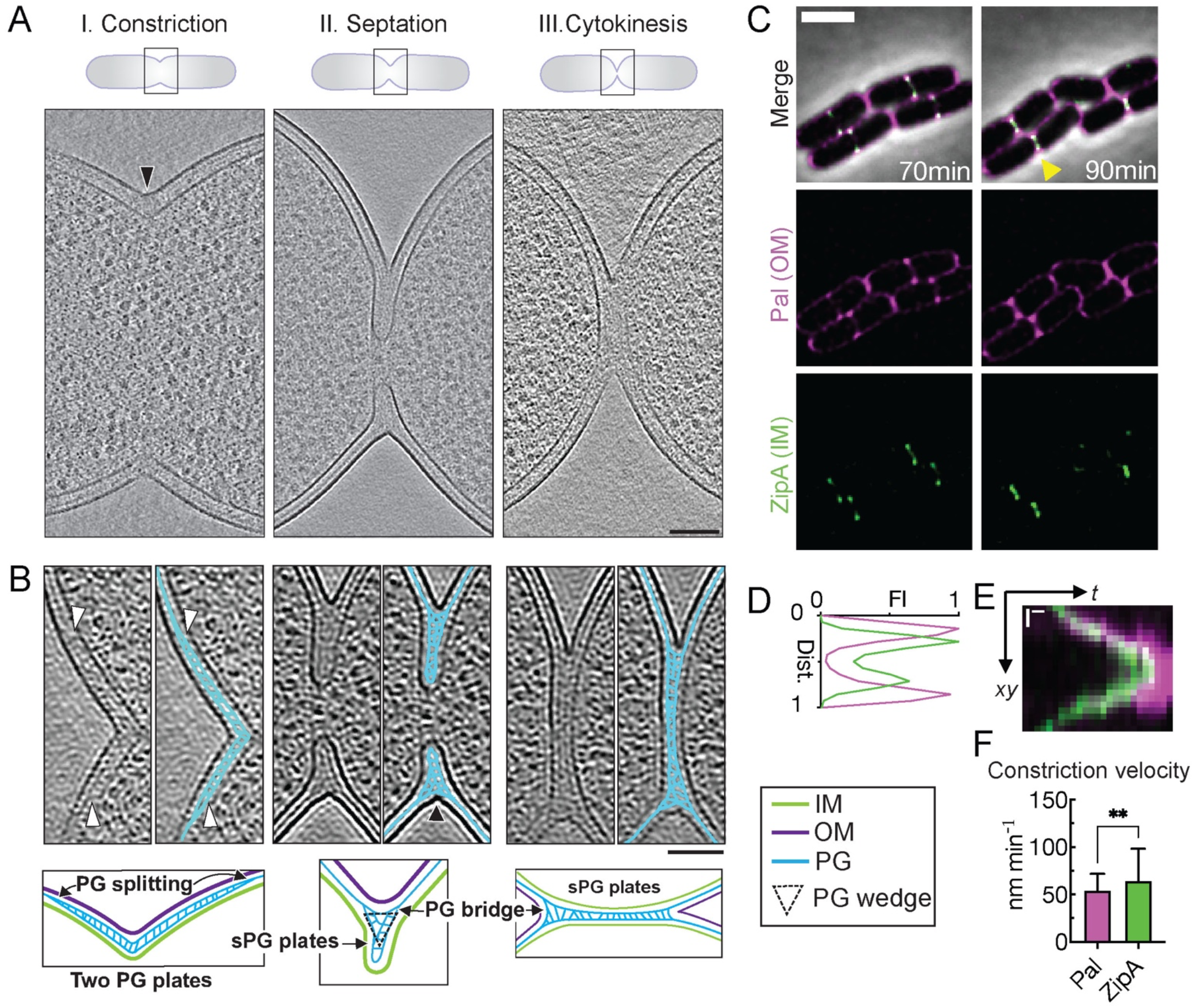
*In situ* cell envelope architecture and dynamics during *E. coli* cell division. (A) Overview of different stages of cell division. Summed, projected central slices of cryo-electron tomograms visualizing different stages in division of wild-type *E. coli* are shown. Black arrowhead indicates division site side displayed in B. (B) NAD-filtered cryo-electron tomograms visualizing the cell wall. Left panels show a 3D slice, right panels show the corresponding slice with segmented PG signal in cyan (see Methods). White arrowheads indicate where the PG layer appears to thicken from one to two layers, and black arrowhead indicates division site side shown in the schematic overview for the septation stage of the visualized cell envelope architecture. (C) Time-lapse series of wild-type *E. coli* expressing Pal-mCherry and ZipA-sfGFP as OM and IM markers, respectively, imaged at 30°C on M9 supplemented with 0.2% casamino acids and D-glucose. Fluorescence signals were deconvolved (see Methods). Yellow triangles mark division sites used for line scans of fluorescence intensity profiles (D) and kymograph analysis of cytokinesis (E). (F) Constriction velocity of IM and OM were derived from the slopes of the fluorescence signals in kymographs (see Methods). All data are expressed as mean + one SD. Two-sided unpaired t-test; ** p < 0.01; N = 150 division kymographs. Scale bar: (A) and (B) = 100 nm, (C) = 2 μm, and (E) = 200 nm (vertical); 5 min (horizontal).

The different division site architectures observed in the cryo-ET analysis suggested that the constriction rate of the IM exceeds that of the OM during division to promote the formation of the partial septa. To investigate this possibility, we followed the constriction dynamics of each membrane using live-cell fluorescence microscopy. The IM was tracked using a superfolder GFP (sfGFP) fusion to the IM-anchored division protein ZipA (ZipA-sfGFP) whereas constriction of the OM was followed using mCherry fused to the OM-localized lipoprotein Pal (Pal-mCherry) (**Fig. 1C and Video S2**). Analyzing the distribution of the fluorescence signals of Pal-mCherry and ZipA-sfGFP at the division site confirmed the further constriction of IM with respect to the OM (**Fig. 1D**) as seen by cryo-ET (**Fig. 1A-B**). Following time-lapse imaging of a large population of cells (N = 150), the invagination rate of each membrane was calculated from kymographs (**Fig. 1E-F**). We found that the IM constricted at a rate of 64.26 ± 33.98 nm/min and the OM at a rate of 55.03 ± 23.25 nm/min. These results are in-line with previously measurements of cytokinesis rates^30,31^. Multiplying the difference in constriction velocity between the membranes by the average time for cytokinesis (16.36 ± 7.49 min) predicts that the IM-IM versus OM-OM diameters will differ by upwards of 147 nm at late stages in cell division, which is in good agreement with our cryo-ET data (**Fig. S2**). Thus, the combined cryo-ET and time-lapse analysis definitively establish that *E. coli* divides by a mixed constriction/septation mechanism. At early stages in the process, invagination of the three envelope layers is well-coordinated, resulting in a V-shaped constriction architecture. However, because the IM invaginates about 9 nm/min more rapidly than the OM, the distance between the two membranes grows as division progresses and a partial septum with a structure of sPG reminiscent of some Gram-positive bacteria is formed. Thus, the division site architectures of diverse bacteria appear to share more similarities than previously anticipated.

### sPG synthesis and remodeling activities of the divisome define division site architecture

We next wanted to determine how the architecture of the division site and the dynamics of its constriction are altered by mutations affecting sPG synthesis and remodeling. The essential PG synthase of the divisome is formed by a complex between FtsW and FtsI (FtsWI) (Taguchi et al., 2019). Following Z-ring assembly at the prospective division site, a regulatory pathway is thought to be initiated that ultimately results in the activation of sPG synthesis by this synthase^34–37^(**Fig. 2A**). Activation is mediated in part via a direct interaction between FtsWI and the FtsQ-FtsL-FtsB (FtsQLB) complex^38^. Although the mechanism is unclear, genetic evidence suggests that the FtsQLB activation event is stimulated by an essential peptide within the division protein FtsN^39^. Another domain of FtsN called the SPOR domain is responsible for concentrating the activation peptide at the division site through its ability to bind sPG that has been processed by PG cleaving enzymes called amidases^39–41^. The amidases remove the stem peptides from the glycan strands as they split the sPG septum to promote OM constriction and daughter cell separation^42^. In the process, they generate the denuded PG glycan strands recognized by the SPOR domain^40,41^. The interplay between the activation of sPG synthesis by FtsN with amidase processing of sPG promoting the recruitment of more FtsN to the division site has been proposed to generate a positive feedback loop, the sPG loop, that drives the activation of sPG biogenesis and cell division^39^. To modulate this process for our analysis, we employed several different mutant strains (**Table S2-S3**): (i) a mutant lacking the SPOR domain of FtsN (*ftsN-ΔSPOR*), (ii) mutants defective for one (*ΔenvC*) or both (*ΔenvC ΔnlpD*) activators required for amidase activity^43^, and (iii) a mutant (*ftsL*\*) encoding a variant of FtsL that hyperactivates sPG synthesis by FtsWI and bypasses the requirement for FtsN^37^ (**Fig. 2A**). A comprehensive gallery of cryo-electron tomograms containing the analyzed *E. coli* strains is provided (**Fig. S3**). A total of 60 tomograms were obtained and analyzed in this study (**Table S1)**.

**Figure 2:**
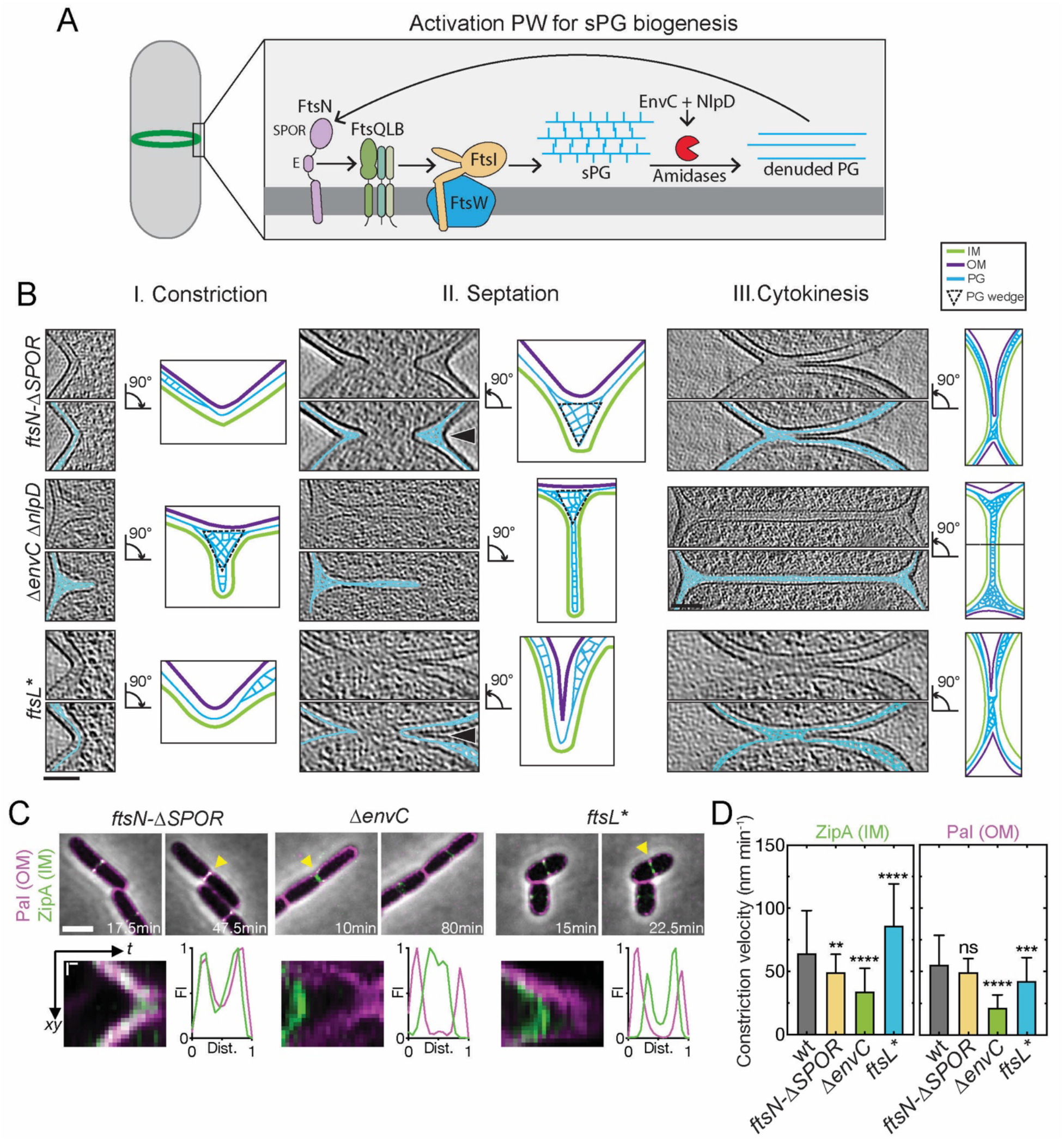
Divisome mutants display altered division site ultrastructure and constriction kinetics. (A) Schematic overview of the septal PG loop pathway for the activation of sPG synthesis (see text for details). (B) NAD-filtered cryo-electron tomograms of division sites in the indicated division mutants of *E. coli*. Top panels show a 3D slice and bottom panels show the corresponding slice with segmented PG signal in cyan (see Methods). Right panels show summary diagrams of the cell envelope architecture visualized by cryo-ET. Black arrowheads indicate division site side represented in the schemes. (C) Time-lapse series of indicated *E. coli* division mutants expressing Pal-mCherry and ZipA-sfGFP as OM and IM markers, respectively, imaged as in Figure 1. Yellow triangles mark division sites used for kymograph analysis and line scans of fluorescence intensity profiles of cytokinesis. (D) Constriction velocity of the IM and OM were determined as in Figure 1. All data are expressed as mean + one SD. Data from wild-type is replotted from Fig. 1F for comparison. Brown-Forsythe and Welch ANOVA tests, differences in significances are tested relative to wild-type; ** p < 0.01, *** = p < 0.001, **** p < 0.0001; N = 150 division kymographs for each strain. Scale bar: (B) = 100 nm and (C) top row = 2 μm and bottom row kymographs = 200 nm (vertical); 5 min (horizontal).

Division sites from cells deleted for the SPOR domain of FtsN resembled those observed previously for *C. crescentus*^16^ (**Fig. 2B**). By cryo-ET, there appeared to be greater coordination between IM and OM constriction throughout the division process with the IM-OM spacing in deeply constricted cells (79.2 ± 21.7 nm) measuring nearly half that of wild-type cells in the septation phase (**Fig. 2B, S2-3, Video S3**). Fluorescence microscopy tracking of IM and OM constriction dynamics confirmed that the rate of constriction for the two membranes was nearly identical in the *ftsN-ΔSPOR* mutant (**Fig. 2C-D, Video S2**). In this case, the rate of IM constriction was reduced compared to wild-type cells (49.56 ± 13.93 versus 64.26 ± 33.98 nm/min) such that it more closely matched the rate of OM constriction **(Fig 2D)**. As a result of the close opposition of the IM and OM, the wedge of sPG observed in filtered tomograms was not as elongated as in wild-type cells such that separate plates of material forming ahead of the wedge could not be clearly visualized (**Fig. 2B**).

In contrast to the coordinated envelope constriction displayed by the *ftsN-ΔSPOR* mutant, cells defective for both amidase activators (*ΔenvC ΔnlpD*) formed a Gram-positive-like septum in which the constriction of the IM was completed without much observable invagination of the OM (IM-OM distance: 296.05 ± 133.45 nm in deeply constricted cells) (**Fig. 2B and S3)**. A similar septal architecture was observed previously for this mutant using conventional EM, but the ultrastructure of the sPG layer was not preserved^44^. NAD-filtering of the current cryo-electron tomograms revealed density corresponding to sPG that was even more clearly discernable as two distinct plates of material than in the partial septa observed in wild-type cells above (**Fig. 1B and 2B**). Furthermore, the tomograms also revealed a triangular wedge of PG material at the outer edges of the septa that was not previously observed in the conventional EM analysis and presumably serves as a roadblock to OM invagination (**Fig. 2B, Video S4**). It was not possible to measure membrane constriction dynamics in live cells of the *ΔenvC ΔnlpD* double mutant due to its poor growth. However, measurements in a mutant lacking only EnvC, the most important of the two amidase activators^43^, revealed a significant disparity between IM and OM constriction rates (**Fig. 2C-D. Video S2**). Even though the IM constriction rate was slower than wild-type or *ftsN-ΔSPOR* cells (33.97 ± 18.42 nm/min) the completion of IM invagination still preceded that of the OM by an average of 36.5 min (**Fig. S4**). Overall, the results with the *ftsN-ΔSPOR* and amidase activation mutants indicate that impairing key components of the divisome involved in sPG synthesis activation and remodeling alters the architecture of the *E. coli* division site such that it begins to resemble that of other distantly related bacteria.

### Hyperactivated sPG biogenesis leads to aberrant division site architecture

To determine the effects of hyperactivated sPG biogenesis on division site architecture, we imaged cells of the *ftsL\** mutant. Strikingly, cryo-ET revealed an altered architecture in which the density corresponding to the wedge of sPG observed in wild-type and other mutant cells was missing, and the envelope at the leading edge of the invagination was 48% thinner than in wild-type (**Fig. 2B and Fig. S5B**). Conversely, the envelope in the nascent polar regions adjacent to the leading edge of the invagination was 15 % thicker in *ftsL\** cells than in wild-type with the bulged areas appearing to contain more PG than normal (**Fig. S5B and Video S5**). As expected from the faster division time measured previously for *ftsL\** mutant^37^, live-cell imaging showed that the constriction velocity of the IM was much greater than for wild-type cells (86.18 ± 33.12 nm versus 64.26 ± 33.98 nm/min) (**Fig. 2C-D, Video S2**). Surprisingly, the rate of OM constriction was measured to be much slower than that of the IM (42.60 ± 18.49 nm versus 55.30 ± 23.25 nm) (**Fig. 2C-D**), a difference that would normally be expected to give rise to cells with partial septa with large distances between the IM and OM. However, partial septa were not observed in the tomograms (**Fig. 2B**). The distances between the IM and OM remained relatively constant in all the cells that were imaged (**Fig. 2B and Fig. S2-3**). This discrepancy is likely due to the PG binding activity of Pal causing the Pal-mCherry OM reporter to get stuck in the thicker PG that accumulates behind the closing septum and therefore to track poorly with the leading edge of the invaginating OM in these cells. In contrast, cryo-ET data directly visualize OM. Notably, upon closer inspection of kymographs of the *ftsL\** mutant, we noticed that one side of the cell constricted much faster than the other (**Fig. 2C**), a phenomenon that was also observed by cryo-ET (**Fig. S6A-C)**. Applying Fourier filtering to kymographs allowed us to distinguish between forward and reverse signal trajectories in order to compare the constriction velocity for each side of the division site and assess the degree of anisotropy during septum closure. We found that *ftsL\** showed a higher but not statistically significant (Kruskal-Wallis ANOVA, p = 0.056) anisotropy score for both IM and OM constriction compared to the other strains (**Fig. S6B-C**). Additionally, when cells were imaged in vertical orientation, the constriction of *ftsL\** cells was observed to be less circular than in wild-type as expected for uneven closure of the division ring (**Fig. S6D**). Thus, circumvention of the normal controls regulating sPG biogenesis in the *ftsL*\* mutant results in aberrant division site geometry and abnormal thickening of the envelope at the cell poles. These cells also lack an observable sPG wedge, which may destabilize the division site and help explain why these mutants were originally found to lyse at elevated temperatures^45,46^.

### sPG degradation activates its synthesis

To better understand the mechanism(s) by which the observed changes in division site architecture are caused by mutations altering divisome components, we measured the rates of sPG synthesis and degradation in *ftsL*, ftsN-ΔSPOR*, and *ΔenvC* cells relative to those of wild-type using two different cytological assays (**Fig. 3 and S7**). The first assay used a pair of compatibly labeled fluorescent D-amino acids (FDAAs), YADA and HADA^47^. In *E. coli*, these probes are primarily incorporated into the peptide stem of PG by alternative crosslinking enzymes called L,D-transpeptidases (LDTs)^48^. Because they do not use the canonical PG synthases (e.g. FtsW, PBP1b, etc.) for incorporation, FDAAs only provide an indirect readout of nascent PG synthesis. Nevertheless, when two probes are used for labeling in sequence, they can accurately label areas of newly synthesized material^49^. In our experiments, we grew cells for an extended period in the presence of YADA to label the entire PG sacculus. A portion of these cultures was then fixed to establish the YADA labeling baseline while other portions were washed, pulsed with HADA for different lengths of time, and then fixed prior to visualization. The intensity of the HADA signal that appeared at midcell after the pulses was used as a measure of the rate of sPG insertion (**Fig. 3A**). Additionally, the YADA label was used to follow the fate of old PG with a comparison of the signal intensity at midcell before and after the HADA pulse providing a measure of sPG degradation (**Fig. 3A**). To measure nascent PG synthesis directly, the second assay employed a MurNAc-alkyne probe^50^ in conjunction with HADA (**Fig. S7**). In this case, cells were pre-labeled with HADA to mark old PG and pulsed with MurNAc-alkyne, which is incorporated into the PG precursor lipid II and used directly by canonical synthases to build new PG^30^. Thus, using click chemistry and a clickable fluorescent dye, the rate of sPG insertion was measured from the MurNAc-alkyne labeling intensity, and comparison of the HADA signal before and after the pulse was used to determine the rate of sPG degradation.

**Figure 3:**
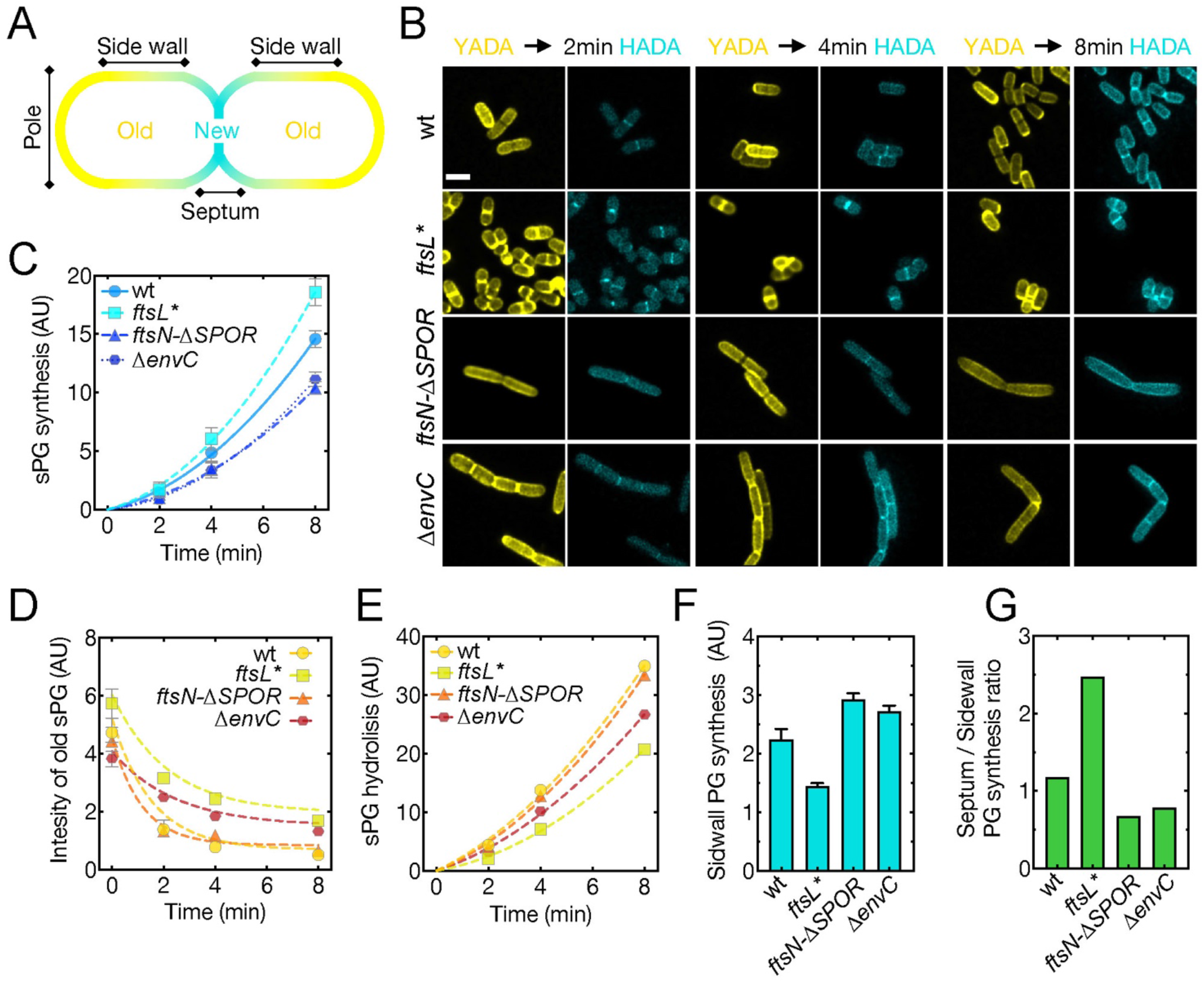
FDAA assay for cell wall synthesis and hydrolysis rates during division and elongation. (A) Schematic representation of labeling patterns observed for the pulse-chase experiment. New cell wall material is labeled with HADA (blue), while old material is stained with YADA (yellow). (B) Representative images of indicated strains after 2, 4 and, 8 min pulses with HADA. Integrated fluorescence intensity was measured and normalized at the division site area for new (C) and old (D) PG. Deriving the integral from these intensity measurements yields the rate of sPG synthesis. Data was fit to a quadratic exponential equation (R-squared > 0.9). (E) Septal PG hydrolysis rates were derived from calculating the reduction in YADA fluorescence intensity over time, as compared to cells fixed prior to chase (time point 0 min, D). (F) Sidewall incorporation of new cell wall material (HADA fluorescence intensity) was measured after 8 min, due low signal intensities in earlier timepoints. (G) Ratio between sPG and sidewall synthesis was calculated by dividing the mean HADA fluorescence intensity for each after the 8 min pulse. All data are represented as mean fluorescence intensity. Error bars represent 95 % confidence intervals. All values are arbitrary fluorescence intensity units and were divided by 1000 for plotting purposes. Scale bar in (A) = 2 μm.

Both PG biogenesis assays yielded qualitatively similar results (**Fig. 3 and S7**), and the measurements of sPG synthesis correlated well with the rates of IM invagination determined above (**Fig. 2D**). The *ftsL\** mutant synthesized sPG faster than all other strains just as it had the fastest rate of IM invagination (**Fig. 2D, 3B, and S7D**). This result confirms that activated FtsQLB complexes indeed hyperactivate sPG synthesis as suggested by their recently reported effects on the dynamic motions displayed by the FtsWI PG synthase^30^. Notably, the dual FDAA assay detected a significant amount of old PG at the division sites prior to the HADA pulse, and this material appeared to be relatively stable during the time course. Additionally, bright foci of old material were also observed at the poles of many cells after extended YADA labeling (**Fig. 3B S8A-B**). This accumulation of old material likely corresponds to the thickened areas of cell wall in the curved regions of septa and nascent poles observed by cryo-ET of the *ftsL\** mutant (**Fig. S5 and S8C-D**), reinforcing the conclusion that short-circuiting the normal controls governing the activation of sPG biogenesis not only leads to more rapid sPG synthesis and septal closure, but also results in the aberrant accumulation of PG within the developing cell poles.

As suggested by their slower than normal rates of IM invagination, both the *ftsN-ΔSPOR*, and *ΔenvC* mutants displayed a reduced rate of sPG synthesis relative to wild-type cells (**Fig. 3C and S7D**). In the MurNAc-alkyne assay, the *ftsN-ΔSPOR* mutant also showed a reduced septal labeling efficiency (**Fig. S7C**), which is another indication of reduced sPG synthesis activity^30^. Although the sPG synthesis rates were similar, the two mutants differed greatly in their rates of sPG degradation. The *ftsN-ΔSPOR* mutant displayed relatively normal rates of sPG degradation whereas *ΔenvC* cells showed reduced turnover of sPG as expected for a mutant lacking an important amidase activator (**Fig. 3D-E and S7E**). The combination of slower sPG synthesis with relatively normal sPG degradation explains the well-coordinated constriction phenotype displayed by the *ftsN-ΔSPOR* mutant in the cryo-ET analysis. Although reduced sPG degradation was expected for the amidase activation mutant based on its cell chaining phenotype and abnormal septal morphology, it has not been directly demonstrated previously. The reduced rate of sPG synthesis in the *ΔenvC* cells is notable, however, because it indicates that proper sPG processing by the amidases is required for normal rates of sPG synthesis. This result along with the reduced rate of sPG synthesis observed for the *ftsN-ΔSPOR* mutant provides strong support for the sPG loop model for PG biogenesis during division, which so far has been based solely on inferences from a genetic analysis of the domain functions of FtsN^39^.

### sPG degradation is required for normal Z-ring formation

In the FDAA labeling experiments we noticed that cells lacking EnvC commonly displayed closely spaced sPG labeling consistent with the aberrant formation of adjacent division sites (**Fig. 4A**). Accordingly, closer examination of the localization of Pal-mCherry in these cells revealed that double bands of the OM marker for sites of cell constriction occurred at a significantly elevated frequency over wild-type or *ftsN-ΔSPOR* cells (**Fig. 4B, E-F**). These double constrictions were typically observed within the cell body, but we also observed constrictions near cell poles, generating what appeared to be minicells (**Fig. 4C, G**). However, free minicells were not observed in the culture, suggesting that these aberrant poles were likely generated from a double constriction event within the cell body, one of which was aborted while the other completed division to leave behind a daughter with a polar constriction. Another defect in division we noticed in the *ΔenvC* cells was the formation of membrane blebs emanating from some of the developing septa (**Fig. 4C**). Cells with such blebs were often observed to lyse, suggesting there was a catastrophic failure in division. Lysed cells and double constrictions were also observed in the cryo-ET analysis of *ΔenvC ΔnlpD* cells (**Fig. 4D**).

**Figure 4:**
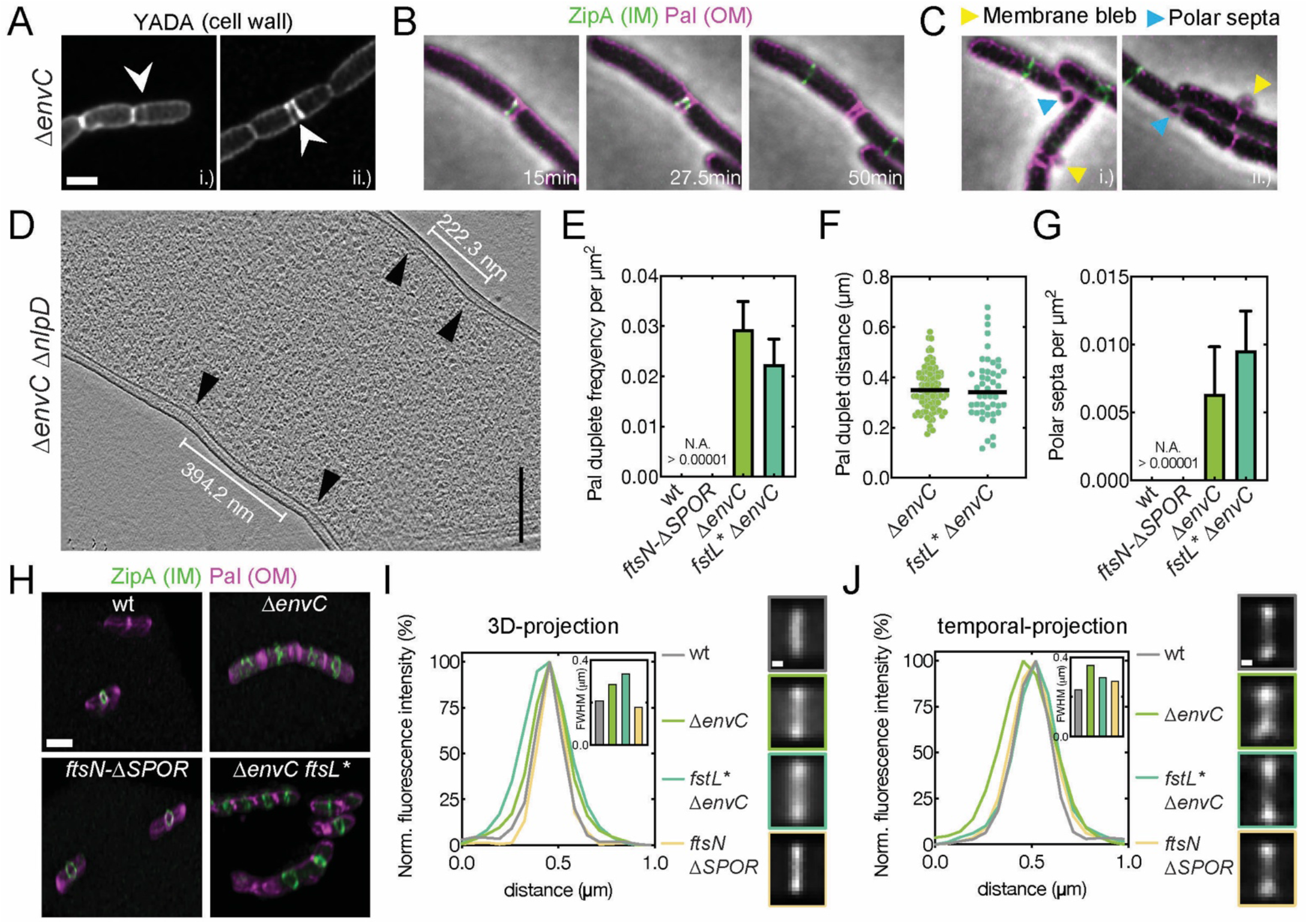
sPG hydrolysis is required for normal Z-ring placement and condensation. (A) Distribution of cell wall material in *ΔenvC* cells was assessed by FDAA staining. Images are sum-projections of a 1 μm spanning z-stack and were deconvolved. White arrowheads indicated double septa. (B) Time-lapse series of a *ΔenvC* mutant expressing Pal-mCherry and ZipA-sfGFP as OM and IM markers, respectively, imaged as in Figure 1. An example of double septum formation is shown. (C) Examples for membrane blebbing (yellow arrowheads) and polar septa (blue arrowheads) formation are highlighted. (D) Formation of double constrictions observed in cryo-electron tomograms of *ΔenvC ΔnlpD* cells. (E) Frequency of double septum formation was quantified from counting the number of Pal-mCherry doublets and normalizing for cell area. No Pal doublets were found > 10,000 cells for wild-type or *ftsN-ΔSPOR* cells. Data is represented as mean + SD. (F) Distance between Pal duplets was measured manually using the line tool in Fiji. (G) Frequency of polar septa was measured for the indicated strains and normalized per cell area. No polar septa were observed in > 10,000 wild-type or *ftsN-ΔSPOR* cells. (H) Three-dimensional maximum intensity renderings showing Z-ring condensation based on ZipA-sfGFP localization. Scale bar = 2 μm. The degree of Z-ring condensation was quantified from averaged fluorescence intensity projections from summed 3D volumes or from 5 time points (corresponding to 10 min) of a time-lapse series (J). Width of the fluorescence signal distribution across the horizontal axis was used to determine degree of condensation. FWHM is shown as an insert. Averaged Z-rings are shown and color-coded according to graphs. Scale bars: (A)-(C) = 2 μm, (D) = 200 nm, (H) = 2 μm, (I)-(J) = 200 nm.

In addition to the aberrant sPG and Pal-mCherry localization, the pattern of ZipA-sfGFP was also altered in the *ΔenvC* mutant relative to wild-type and the other mutants (**Fig. 4H, Video S6**). ZipA is an FtsZ-binding protein and a marker for the Z-ring^51^. Many ZipA-sfGFP structures in cells lacking EnvC were diffuse, suggesting that they are having difficulty condensing into the tight Z-ring structure typical of normal cells (**Fig. 4H**). Consistent with this possibility, averaging the ZipA-sfGFP signals for a population of cells (N = 100) either over a 10-minute time window or over a 2 μm volume spanning midcell showed that the fluorescent signal was more broadly distributed in the *ΔenvC* mutant than in wild-type cells or cells of the other mutants (**Fig. 4I-J**). Notably, this phenotype was not suppressed by combining the sPG synthesis activating *ftsL*\* mutation with *ΔenvC*, indicating that it likely stems from the loss of sPG processing not its collateral effect on sPG synthesis (**Fig. 4H-J**). A similar diffusion of the Z-ring signal has been observed for cells defective for FtsZ-binding proteins like ZapA that are thought to bundle FtsZ polymers to condense the ring^52,53^. These results therefore suggest a previously unappreciated role for sPG hydrolysis by the amidases in the periplasm in promoting Z-ring condensation in the cytoplasm. The observation of double septa and failed division sites also indicates an unexpected role for amidase activation in division site stability and/or placement in addition to their known function in cell separation (see Discussion).

### Competition between elongation and sPG biogenesis determines polar shape

The idea that the cell elongation and division machineries might be in competition with one another was proposed many years ago^54,55^. However, experimental support for such a competition has been limited, and it has never been demonstrated that competition occurs at the level of PG biogenesis. To investigate this possibility further, we took advantage of the PG labeling assays used to measure sPG biogenesis to also quantify the incorporation of new PG material into the sidewall (cell cylinder) during the MurNAc-alkyne or HADA pulses (**Fig. 3F and S7F**). Importantly, we found that sidewall PG and sPG synthesis were inversely proportional in the different strains we studied. Sidewall PG incorporation was highest in the *ftsN-ΔSPOR* mutant, which had one of the lowest rates of sPG synthesis (**Fig. 3 and S7**). Conversely, sidewall PG synthesis was the lowest in the *ftsL*\* mutant that displayed the highest measured rates of sPG production (**Fig. 3 and S7**). In addition to the rates of sidewall PG biogenesis being inversely proportional to that of sPG biogenesis, they were also found to be directly related to the single cell elongation rates of the division mutants, with *ftsN-ΔSPOR* mutants elongating 1.5 times faster than *ftsL\** cells (**Fig. 5A**). Another measure of cell elongation activity is the circumferential motions of MreB and other components of the Rod complex around the cell cylinder^56–58^. We therefore tracked the motion of a mNeonGreen fusion to MreB in wild-type and mutant cells using a combination of structured Illumination microscopy and total internal reflection fluorescence (SIM-TIRF) imaging. Consistent with the sPG synthesis measurements, the total number of directionally moving MreB filaments per area was significantly reduced in *ftsL\** and *ΔenvC* cells (**Fig. 5B-C, Video S7**). Interestingly, while loss-of-function mutations in the division machinery (e.g., *ftsN-ΔSPOR, ΔenvC*) result in an increased cell length, hyperactivation of the division machinery (e.g., *ftsL\**) leads to shorter cells than wt (**Fig. S9A**). Concomitantly, mutants displaying reduced sidewall PG incorporation rates and fewer directionally moving MreB filaments such as *ftsL\** and *ΔenvC*, were also found to display significantly increased cell width (**Fig. S9B**), which are indicative of reduced Rod complex activity^59^.

**Figure 5:**
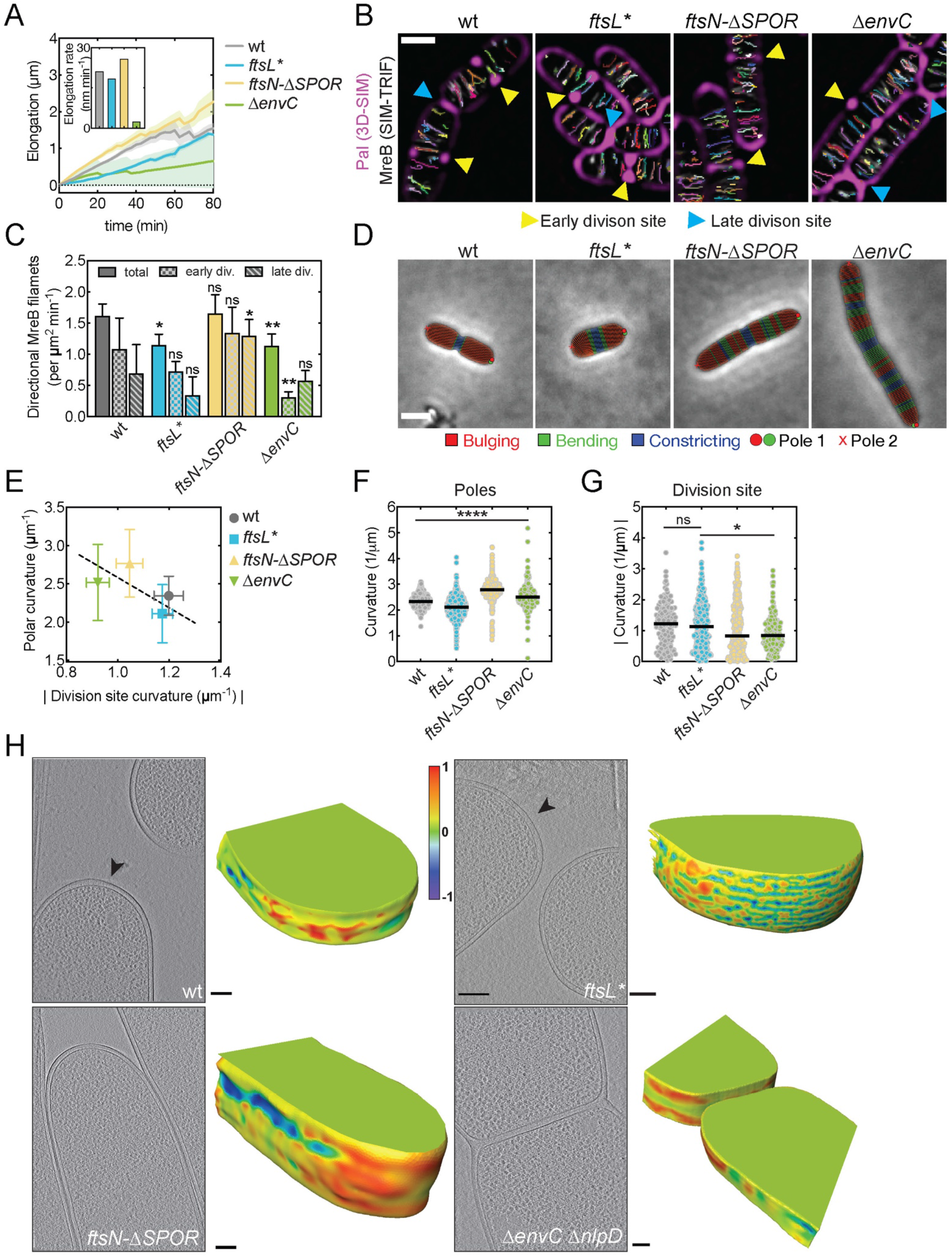
Competition between the divisome and elongation machinery defines polar cell shape. (A) Single-cell elongation was measured for indicated mutants grown at 30°C on M9 supplemented with 0.2% casamino acids and D-glucose for 80 min. Elongation rate (insert) was determined from the slope of a best fit linear regression model. (B) MreB dynamics were followed using an mNeonGreen sandwich fusion (MreB^sw-mNG^) in the indicated strains for 3 min at 3 second acquisition frame rate using SIM-TIRF microscopy. Time-lapse series were sum projected and overlayed with single particle tracking results from TrackMate and 3D-SIM Pal-mCherry reference images. Pal-mCherry signal serves to identify constricted cells. Early division site (yellow arrowheads) displayed Pal foci which were resolvable as two distinct foci, whereas late division sites (blue arrowheads) displayed a continuous Pal signal across the cell body, indicative of complete or near complete cytokinesis. (C) Directionally moving MreB tracks were filtered by MSD analysis (see Methods) and plotted as mean + SD and normalized by cell area. One-way ANOVA test, differences in significances are tested relative to wild-type for each corresponding stage; * p <0.05, ** p < 0.01, ns = non-significant. (D) Representative phase contrast micrographs showing segmented cells in *Morphometrics* for the indicated division mutants. Color-coded legend indicates cell regions with positive curvature on both sides (red, bulging), negative curvature on both sides (blue, constricting) and positive curvature on one side while negative curvature on the other side (green, bending). Red dots indicate poles. Scale bar = 2 μm. (E-G) Polar curvature was measured by the two highest points of positive cell outline curvature, while constriction curvature was assessed by measuring the opposing contour-matched lowest curvature values at the division site using *Morphometrics* (see Methods). Polar and division site curvatures are negatively correlated. Ordinary One-way ANOVA; * p <0.05, **** p < 0.0001, ns = non-significant. (H) Summed, projected central 3D slices through cryo-electron tomograms visualizing cell poles. Black arrowheads indicate 3D rendered pole. The corresponding 3D-volume renderings show the type of curvature present on the pole surface determined by shape index (see Methods). Summed projection images are scaled. Scale bars: (B) = 1 μm, (D) = 2 μm (H) summed projection images = 200 nm and 3D renderings = 100 nm. 011

Notably, the observed interplay between cell elongation and division appeared to impact the geometry of the division site and the shape of the daughter cell poles (**Fig. 5D**). The *ftsN-ΔSPOR* mutant which elongates more rapidly and constricts slower, displayed an elongated division site area and a shallower OM invagination angle at midcell as compared to wild-type cells (**Fig. 5D-G**). This altered constriction geometry was also observable in the cryo-electron tomograms of dividing cells and correspondingly gave rise to daughter cells with pointier poles than wild-type (**Fig. 5F, H**). On the other hand, the rapidly constricting *ftsL\** mutant formed daughters with relatively blunt cell poles (**Fig. 5F, H**).

We reasoned that the variation in division site and polar geometry among the different strains could be related to the activity of the Rod complex at or near the division site. In this case, mutants that take longer to complete the division process (such as *ftsN-ΔSPOR*) would allow for more elongation activity in the curved cell region adjacent to the site of constriction, thereby generating shallower constrictions and elongated poles (**Fig. 5E-G**). Conversely, mutants that divide rapidly (such as *ftsL\**) would have less opportunity to elongate the region around the division site and thus form steep constrictions and blunt poles (**Fig. 5E-G**). To investigate this possibility, we quantified the number of directionally moving MreB filaments in proximity (≤ 200 nm) to cell constrictions highlighted by Pal-mCherry foci at mid-cell (**Fig. 5B-C, Video S7**). Directionally moving MreB filaments were readily observed to pass through division sites in both early and late pre-divisional cells in all strains tested. Notably, however, the *ftsN-ΔSPOR* mutant displayed more MreB tracks at the division site at late stages of division than all other strains, and the *ftsL\** mutant showed the least number of total MreB tracks at the division site (**Fig. 5C**). Thus, the density of MreB tracks at the division site for these cells correlates well with the degree of constriction site and cell pole elongation observed for the different strains. The outlier was the *ΔenvC* mutant, which had an inverted trend of having fewer directionally moving MreB tracks at early division stages than at later points (**Fig. 5C**). We suspect that this change is due to the defect in sPG splitting, which causes a steep curvature of the inner membrane at early points in division that is likely to be unfavorable for MreB localization^60,61^. However, at later stages when sPG processing eventually allows for slow constriction of the OM, this curvature likely becomes more favorable for MreB localization allowing elongation to occur near the division site to generate a shallow constriction like that of the *ftsN-ΔSPOR* mutant. Overall, these results not only provide strong support for a competition between the PG biosynthetic machineries involved in cell elongation and division, but they also highlight the potential for this competition to define the morphology of the daughter cell poles.

## Discussion

Understanding the mechanisms regulating bacterial division requires unambiguous visualization of the native cell envelope architecture at division sites. Previously obtained data from EM^28,62^, CEMOVIS^63,64^ and cryo-ET^17,65^ of Gram-negative bacteria, lacked a clear view of the sPG material produced by the division machinery. Here, we overcome this limitation by combining cryo-FIB milling with cryo-ET to visualize the division site of *E. coli*, providing 3D visualization of the *in situ* division site architecture at nanometer resolution. Our data reveal several new ultrastructural features of the Gram-negative cell wall, including a wedge-like formation of sPG that may play an important role in fortifying the septum against osmotic rupture. Integrating these observations with dynamic measurements of sPG synthesis and remodeling in several division mutants led to several new mechanistic insights into the process of cell division and how its interplay with cell elongation can modulate cell shape (**Fig. 6**).

**Figure 6:**
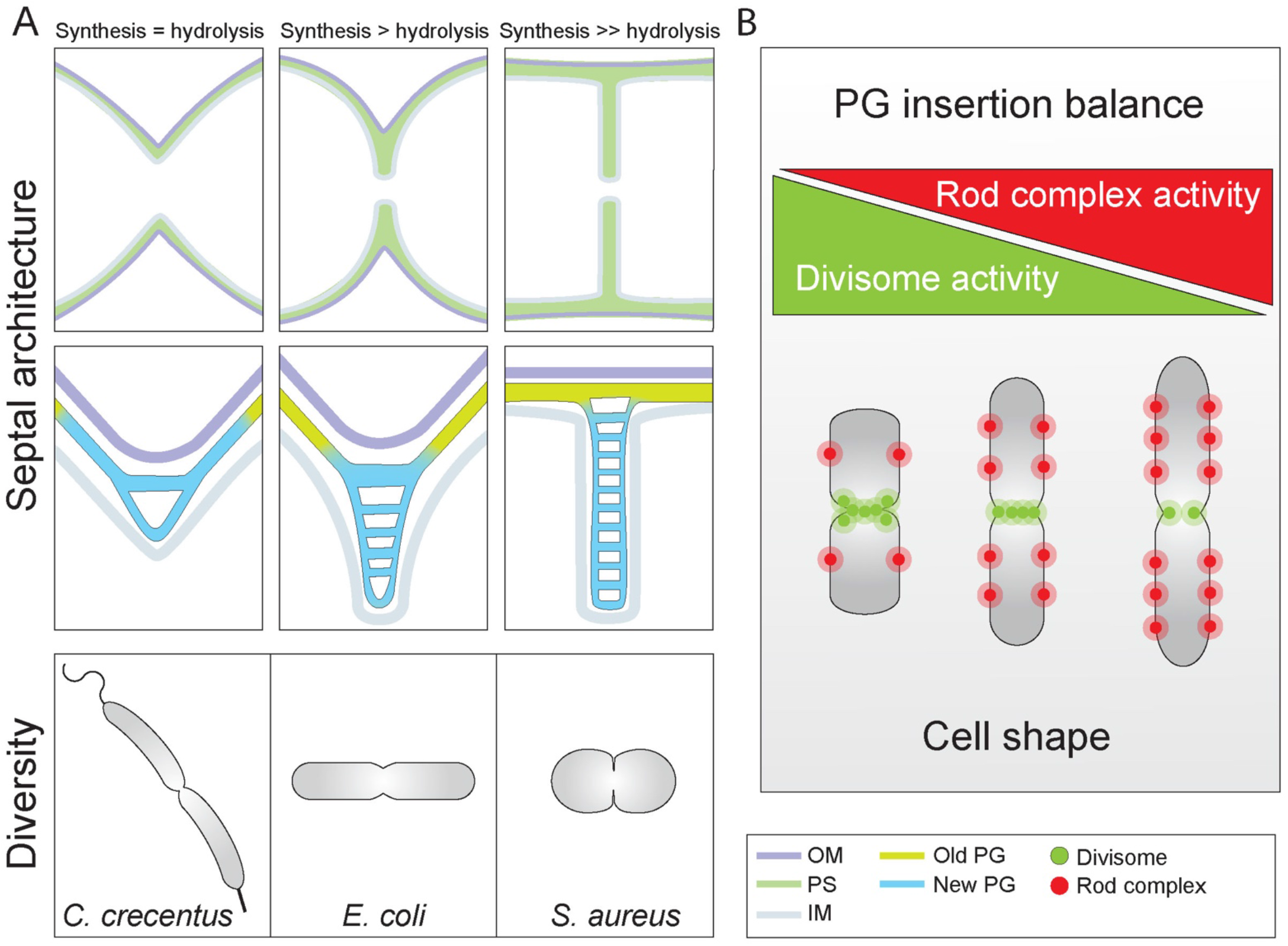
Septal PG architecture and divisome activity modulate bacterial morphogenesis. (A) A wide range of different sPG architectures can result from the same division machinery through altering rates of cell wall synthesis and hydrolysis. Constrictive mode of cell division where OM and IM invaginate at similar velocities, a phenotype commonly associated with *C. crescentus*, is the result of lower sPG synthesis rates. Here, the septal cell wall displays a V-shaped wedge where no sPG plates are present. In contrast, inhibition of sPG hydrolysis causes a temporal separation of IM and OM constriction leading to septation. These septa are reminiscent to those of Gram-positive bacteria such as *S. aureus*, displaying two distinctive plates of sPG linked together by a PG bridge. Wild-type *E. coli* displays a mixed constriction-septation phenotype resulting in a partial septum at later stages of the division process. (B) The activities of the two major synthetic cell wall machineries, the Rod complex and the divisome, are anticorrelated, likely due to competition for limited substrate (lipid II). Balance of their relative activities determines the shape of the cell division site and the resulting poles they form. Cells with higher Rod complex activity are thinner and form pointier poles, while cells with elevated divisome activity are shorter and wider with blunt poles.

### Architecture of the sPG layer

In tomograms of wild-type cells just starting to constrict, all three envelope layers appeared to be invaginating in concert, and little change in the sPG relative to the sidewall PG was evident. However, cells in advanced stages of division had an increased distance between the IM and OM and formed a partial septum. In NAD-filtered tomograms of these cells, a triangular wedge of what is likely to be sPG is observed at the lagging edge of the septum closest to the tip of the invaginating OM. The wedge thins as it approaches the leading edge of the closing IM, and in this narrow portion of the septum, two dense tracks of material are often discernable that correspond to the PG layers that will eventually fortify the daughter cell poles. In cells defective for sPG processing by the amidases, the sPG wedge structure is more prominent than in wild-type cells and it appears to impede the invagination of the OM. We thus infer that amidases process this structure to allow constriction of the OM. Furthermore, because the sPG wedge is observed in deeply constricted wild-type cells as well as unconstricted amidase activation mutants, we suspect that the structure is dynamic with its lagging edge being degraded as new wedge material is deposited at the leading edge. Such a spatial separation of synthesis and degradation would allow the sPG wedge to move in a treadmill-like fashion ahead of the OM as the septum closes.

Premature separation of the daughter cell PG layers before a continuous wall structure is formed between them would create a tear in the PG matrix and ultimately cause osmotic cell lysis. The positioning of the sPG wedge at the lagging edge of the septum suggests that it might function to prevent such a catastrophe by providing a buffer of extra PG material linking the daughter cells. Accordingly, the *ftsL*\* mutant lacks an observable sPG wedge structure and has been shown to have a temperature-sensitive lysis phenotype^45,46^. We therefore conclude that the observed wedge structure plays a role in septal integrity similar to the bridging portion of the π-like PG structure observed in dividing *S. aureus* cells^10^. However, the difference is that the wedge is likely to be continuously remodeled as cells divide gradually whereas the bridge in *S. aureus* appears to provide a static connection between daughter cells until the end of the division process when lesions in the bridge are created and crack propagation results in extremely rapid cell separation^66^.

The enzymes responsible for creating the sPG wedge remain to be identified, but our results with the *ftsL*\* mutant suggest that it is not made by FtsWI synthase. This mutant is thought to hyperactivate FtsWI^30,35– 38^. Therefore, if the wedge were produced by the FtsWI synthase, the *ftsL*\* mutant would be expected to produce a thicker or otherwise larger wedge. Instead, it lacks a wedge altogether, suggesting that enhanced FtsWI activity disrupts biogenesis of the sPG wedge by other synthases. An attractive candidate for this additional synthase is the class A penicillin-binding protein (aPBP) PBP1b. Inactivation of PBP1b has been found to be synthetically lethal with defects in FtsWI activation, and the affected mutants were found to lyse due to septal lesions, suggesting that this aPBP promotes division site stability^35,67^. The location of the wedge at the lagging edge of the division site closest to the OM is also consistent with a role for PBP1b in its construction given that this enzyme and related aPBPs require activation by lipoproteins anchored in the OM to make PG^68,69^. Thus, the outer fork of the division site where the wedge is located is the only place where aPBPs would be predicted to be functional. Although further work will be required to test this model, it provides an attractive explanation for the division of labor between the aPBP and FtsWI synthases at the site of cell constriction, with the FtsWI synthase promoting ingrowth of the PG layer and the aPBPs providing backfill to counter the activity of sPG remodeling enzymes, thereby stabilizing the septum and preventing lysis.

### The sPG activation loop

The proposal that FtsN and the amidase enzymes cooperate in a positive feedback loop that promotes sPG synthesis is based on insightful reasoning aimed at explaining how the different domains of FtsN might work together^39^. However, this model lacks direct experimental support. A major prediction of the model is that proper sPG synthesis should require both the SPOR domain of FtsN to recognize denuded glycan strands at the division site and amidase activity to generate them^39^. Our results indicate that this is indeed the case. Both the *ftsN-ΔSPOR* mutant and a mutant defective for amidase activation were found to synthesize sPG at a reduced rate compared to wild-type cells. Notably, sPG is still made in these mutants, and even cells defective for all amidases or amidase activators can complete septum formation in *E. coli*. In the case of the *ftsN-ΔSPOR* mutant, other SPOR domain containing division proteins like DedD are likely to be promoting feedback loops analogous to that of FtsN^70^. However, the observation that mutants defective for all cell division amidases still make septa suggests that although efficient sPG biogenesis depends on the sPG loop, it is not absolutely required to make sPG.

In addition to stimulating sPG synthesis, the sPG activation pathway in which the sPG loop operates also appears to be important for normal septal architecture. The *ftsL*\* mutant hyperactivates the FtsWI synthase and eliminates the strict FtsN requirement for sPG biogenesis^37^. This short-circuiting of the normal division activation pathway not only causes the loss of the sPG wedge structure observed in wild-type and other mutant cells, but it also promotes the aberrant accumulation of PG within the developing cell poles. Whether this accumulation results from inappropriate activation of PG synthesis by FtsWI or PBP1b, the improper turnover of the deposited material, or some combination of the two remains unknown. Nevertheless, what is clear is that bypassing the normal controls involved in sPG activation and speeding up the process has adverse consequences on the envelope architecture of the poles that are formed. We therefore infer that the normal divisome activation pathway serves an important function in coordinating different activities of the machinery to ensure that division is successfully completed once it is initiated and that the polar end products it produces have a uniform surface.

### PG hydrolysis and the Z-ring

Over the years, many factors have been identified that help to position the Z-ring and condense it at midcell. These proteins are typically thought to act upstream of divisome maturation and the activation of sPG biogenesis^6^. However, our results have uncovered an unexpected connection between the activation of sPG processing by the amidase and Z-ring structure, suggesting there is feedback to the Z-ring from events downstream of sPG synthesis activation. Z-rings were found to be poorly condensed in mutant cells lacking the amidase activator EnvC. Additionally, closely spaced constrictions or areas of sPG biogenesis were also observed at an elevated frequency in these cells, suggesting that division sites are unstable and fail before they complete the division process. Notably, closely spaced constrictions were also evident in prior EM of chaining cells lacking all amidases^42^, and amidase activity has been shown to be critical for the completion of cell constriction in the related Gram-negative bacterium *Pseudomonas aeruginosa*^71^. Taken together, these results suggest the counterintuitive notion that sPG degradation by the amidases is required to stabilize the divisome, most likely via a positive influence on the structure of the Z-ring that promotes or maintains its condensation. Given that the amidases act on sPG in the periplasm, they are unlikely to directly modulate FtsZ activity. Rather, their effect is probably mediated through SPOR domain proteins like FtsN and DedD that bind the amidase processed glycans^40,41^. These proteins have transmembrane domains and N-terminal cytoplasmic tails, which in the case of FtsN is known to associate with the FtsZ-binding protein FtsA^72,73^. Thus, the status of sPG biogenesis in the periplasm could be communicated to the Z-ring in the cytoplasm using the binding of SPOR domain proteins to denuded glycans as a proxy. Whether the effect might be mediated simply by concentrating the cytoplasmic domains of SPOR proteins at the division site to modulate the activity of FtsZ-binding proteins or via more complex mechanisms requires further investigation, but the general picture that emerges is that the divisome Nactivation pathway is not a one-way street from Z-ring formation to sPG synthesis and processing. The Z-ring is also likely to be receiving stabilizing/activating signals back from the PG biogenesis machinery.

### Cell shape and the balance between cell division and elongation

The idea that the cell elongation and division machineries may be in competition with one another has been discussed in the field for some time^54,55^. However, it has only been recently that evidence for such a completion has been presented. Cells with a temperature-sensitive allele of *ftsI* (*ftsI*23) were found to elongate faster than wild-type during growth at the permissive temperature^74^. Conversely, hyperactivated for the Rod complex were found to be longer than a wild-type control, suggesting that division occurs less frequently when cell elongation is stimulated^75^. Finally, the overproduction of cell wall endopeptidases implicated in cell elongation was found to cause lethal cell division defects in mutants impaired for FtsWI activation^67^. Although these findings are supportive of cell elongation occurring at the expense of division and vice versa, it has remained unclear whether this competition occurs at the level of PG synthesis. Here, we used three independent assays (NAM/FDAA incorporation, single-cell elongation rate, and MreB tracking) to demonstrate that septal and side wall PG synthesis rates are inversely correlated to each other, providing strong support for antagonism between the activities of the elongation and division systems which most likely stems from a competition for the limited supply of the lipid II PG precursor. Importantly, our results indicate that this competition does not just affect cell width or how long or short cells are. It also influences the geometry of the septum and the shape of the daughter cell poles. Thus, modulation of the relative activities of the elongation and division systems is likely to play an important role in the generating the diversity of shapes observed among different bacteria.

## Acknowledgments

We are grateful to Phat Vinh Dip, Edward Brignole and Anna Osherov at the MIT.nano cryo-EM facility and KangKang Song and Chen Xu at the University of Massachusetts cryo-EM facility for providing access to the cryo-EM microscopes and for all their help, advice, and maintenance of cryo-EM equipment. Furthermore, we would like to express our gratitude to the MicRoN imaging core at Harvard Medical School, for excellent advice on live cell imaging and maintenance of fluorescence microscopes. We would also like to thank Calixto Saenz and the Microfabrication Core facility at the department of Systems Biology at Harvard Medical School, for the micro-pillar design, fabrication, and consultations. Lastly, we are grateful to Thomas Bartlett and Erkin Kuru for advice and helpful discussions on FDAA and NAM labeling experiments. P.P.N. was supported by the Swiss National Science Foundation (SNF) with both Early Postdoc.Mobility P2BSP3_188112 and Postdoc.Mobility fellowships P400PB_199252. A.V. was supported by a EMBO long-term postdoctoral fellowship ALTF_89-2019 and a SNF Postdoc.Mobility fellowship P500PB_203143. C.A. is funded by Charles University with a PRIMUS grant (PRIMUS/20/SCI/015). This work was also supported by funding from the National Institutes of Health (R35GM142553 to L.H.C. and R01AI083365 to T.G.B.) and Investigator funds from the Howard Hughes Medical Institute (T.G.B.).

## Author Contributions

P.P.N and A.V. conceived the project, performed experiments, and analyzed and interpreted the data. P.P.N performed cryo-FIB / cryo-ET and image processing. A.V. generated mutants and performed fluorescence microscopy experiments. V.Y.A. performed 3D segmentations of cryo-ET data. P.M.L. established SIM-TIRF workflow and assisted in data collection. C.A. contributed to cell morphology analyses. L.H.C and T.G.B provided infrastructure and scientific advice. P.P.N, A.V., L.H.C. and T.G.B. wrote the manuscript with input from all authors.

## METHODS

### Media, bacterial strains, and mutagenesis

Indicated strain derivatives of *E. coli* MG1655 used in this study are listed in Table S2-S3. Bacteria were grown in LB (1% Tryptone, 0.5% yeast extract, 0.5% NaCl) or M9 medium^76^ supplemented with 0.2% D-glucose and casamino acids. For selection, antibiotics were used at 10 μg ml^-1^ (tetracycline), 25 μg ml^-1^ (chloramphenicol), and 50 (kanamycin, ampicillin) μg ml^-1^. Mutant alleles were moved between strains using phage P1 transduction. If necessary, the antibiotic cassette was removed using FLP recombinase expressed from pCP20^77^. All mutagenesis procedures were confirmed by PCR.

### Cryo-EM specimen preparation

Figure S1 summarizes the cryo-FIB / cryo-ET pipeline utilized in this study. Bacterial strains were grown overnight in LB media, back diluted 1:1000 and incubated shaking at 37°C, 250 rpm to OD_600_ = 0.3. Cells were harvested by centrifugation (2 min, 5000 x g, RT) and resuspended in LB media to a final OD_600_ = 0.6. Three μL of cell culture were applied to Cflat-2/1 200 mesh copper or gold grids (Electron Microscopy Sciences) glow discharged for 30 seconds at 15 mA. Grids were plunge-frozen in liquid ethane^78^ with a FEI Vitrobot Mark IV (Thermo Fisher Scientific) at RT, 100% humidity with a waiting time of 10 seconds, one-side blotting time of 13 seconds and blotting force of 10. Customized parafilm sheets were used for one-side blotting. All subsequent grid handling and transfers were performed in liquid nitrogen. Grids were clipped onto cryo-FIB autogrids (Thermo Fisher Scientific).

### Cryo-FIB milling

Grids were loaded in Aquilos 2 Cryo-FIB (Thermo Fisher Scientific). Specimen was sputter coated inside the cryo-FIB chamber with inorganic platinum, and an integrated gas injection system (GIS) was used to deposit an organometallic platinum layer to protect the specimen surface and avoid uneven thinning of cells. Cryo-FIB milling was performed on the specimen using two rectangular patterns to mill top and bottom parts of cells, and two extra rectangular patterns were used to create micro-expansion joints to improve lamellae instability^79^. Cryo-FIB milling was performed at a nominal tilt angle of 14°-18° which translates into a milling angle of 7°-11°^80^. Cryo-FIB milling was performed in several steps of decreasing ion beam currents ranging from 0.5 nA to 10 pA and decreasing thickness to obtain 100-200 nm lamellae.

### Cryo-electron tomography

All imaging was done on a FEI Titan Krios (Thermo Fisher Scientific) transmission electron microscope operated at 300KeV equipped with a Gatan BioQuantum K3 energy filter (20 eV zero-loss filtering) and a Gatan K3 direct electron detector. Prior to data acquisition, a full K3 gain reference was acquired, and ZLP and BioQuantum energy filter were finely tuned. The nominal magnification for data collection was of 42,000x or 33,000x, giving a calibrated 4K pixel size of 2.193 Å and 2.565/2.758 Å, respectively. Data collection was performed in the nanoprobe mode using the SerialEM^81^ or Thermo Scientific Tomography 5.3 software. The tilt range varied depending on the lamella, but generally was from -70° to 70° in 2° steps following the dose-symmetric tilt scheme^82^. Tilt images were acquired as 8K x 11K super-resolution movies of 4-8 frames with a set dose rate of 1.5-3 e-/Å/sec. Tilt series were collected at a range of nominal defoci between -3.5 and -5.0 μm and a target total dose of 80 to 180 e^−^/Å^2^ (Table S1).

### Cryo-electron tomography image processing

Acquired tilted super-resolution movies were motion corrected and Fourier cropped to 4K x 5K stacks, minimizing aliasing effects using *framealign* from IMOD^83^. Tilt series were aligned using *etomo* in IMOD^84^ and *Dynamo*. CTF-estimation was performed in IMOD and/or using customized MATLAB scripts. CTF-correction was performed by *ctfphaseflip* program in IMOD^85^. CTF-corrected unbinned tomograms were reconstructed by weighted back projection with and without a SIRT-like filter and subsequently 2x, 4x and 8x binned in IMOD^84^.

Bandpass filtering and summed projection of cryo-tomogram slices was performed in *Dynamo*^86–89^ complemented with customized MATLAB scripts. Gaussian and NAD-filtering were performed in Amira (Thermo Fisher Scientific) for visualization purposes. NAD-filtering was applied using the command ‘*Anisotropic Diffusion*’ in 3D mode for 5 iterations. Gaussian filtering was done by applying the command ‘*Gaussian Filter*’ under 3D mode with a Kernel size factor of 3. Whole 3D volume FFT filtering was performed in IMOD.

Three-dimensional pole curvature rendering was performed in Amira by applying the commands ‘*Curvature*’ based on the triangulated 3D mesh and ‘*Shape Index*’ as implemented in Amira^90^. Shape index (SI) computes the surface scalar field which values are equal to

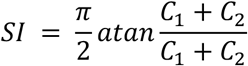

where *C*_1_ and *C*_2_ are the two principal curvatures. Shape index ranges from -1 to 1, negative values indicate negative curvature, positive values indicate positive curvature and values close to 0 indicates flatness of the surface ^90^ (Fig. 5).

### Quantification of cryo-ET data

#### Division site dimensions

Summed projection images of cryo-ET tomograms were used to measure cell dimensions at the division site. Measurements were performed in Fiji^91^ using the ‘point to point’ measuring tool. Measurements were from IM to IM and from OM-OM. Descriptive statistics indicating N and mean ± SEM can be found in Fig. S2.

#### Periplasmic space

Measurements of periplasmic space thickness were performed from OM to IM in the cell areas referred to here as ‘side wall’, ‘pole’ and ‘curve’. Measurements from IM to IM were performed in the cell area defined in this study as septum (S5 and S8). We used a customized macro in Fiji that measures thirty Euclidean distances from surface-to-surface areas in nm, e.g., from IM to IM at the septum. For these thirty single measurements the mean was calculated, yielding a final single mean value per defined subcellular localization, e.g., septum.

### Sample preparation for live cell imaging

Overnight cultures of indicated *E. coli* strains were grown in LB supplemented with appropriate antibiotics at 37°C. The next day, cells were harvested by centrifugation (2 min, 5000 x g, RT) and washed 2x with M9 medium. Day cultures were back diluted (1:1000) and grown in M9 (0.2 % D-glucose, casamino acids) supplemented with 50 μM IPTG and appropriate antibiotics at 30°C until OD_600_ = 0.2 - 0.4. Cells were harvested (2 min, 5000 x g, RT) and resuspended in 1/10^th^ of the original volume. Two microliters of this cell suspension were added onto a 1 % (w/v) agarose in M9 (0.2 % D-glucose, casamino acids) pad supplemented with 50 μM IPTG and covered with a #1.5 coverslip.

### Live cell imaging

All samples were imaged on a Nikon Ti-E inverted widefield microscope equipped with a fully motorized stage and perfect focus system. Images were acquired using a 1.45 NA Plan Apo 100x Ph3 DM objective lens with Cargille Type 37 immersion oil. Fluorescence was excited using a Lumencore SpectraX LED light engine and filtered using ET-GFP (Chroma #49002) and ET-mCherry (Chroma #49008) filter sets. Images were recorded on an Andor Zyla 4.2 Plus sCMOS camera (65 nm pixel size) using Nikon Elements (v5.10) acquisition software. For subsequent deconvolution procedures, three 200 nm spaced Z-planes were acquired for both fluorescence channels using 100% LED output power and 50 ms exposure. Temperature was maintained at 30°C using a custom-made environmental enclosure. After a 20 min acclimatization period, cells were imaged at a 2.5 min acquisition frame rate for a total observation time of 1-4 h.

### Image processing for fluorescence microscopy

First, time-lapse series and Z-stacks were drift corrected using a customized StackReg plugin in Fiji ^91,92^. Subsequently, fluorescence images were deconvolved using the classical maximum likelihood estimation (CMLE) algorithm in Huygens Essential v19.10 (SVI) employing an experimentally derived PSF from 100 nm TetraSpeck beads (Thermofisher). Image reconstruction was performed over 50 iterations with a quality threshold of 0.01 and a signal-to-noise ratio set to 20 for live-cell imaging and 40 for fluorescent cell wall probes in fixed samples. Chromatic aberrations between different fluorescent wavelengths were post-corrected using the Chromatic Aberration Corrector in Huygens from TetraSpeck bead template. The same image reconstruction parameters and chromatic aberration templates were applied to images which were compared to each other. Last, reconstructed fluorescence images were merged back to phase contrast images and rendered for figure or movie display with Fiji.

### Measuring cell envelope constriction dynamics

Fluorescent fusions to IM anchored protein ZipA, and OM lipoprotein Pal, allowed us to determine the respective position of the different cell envelope layers during division. Constriction dynamics of IM and OM were derived from kymographs generated using the Fiji plugin *KymographClear*^93^ and automictically split into forward and reverse trajectories using Fourier filtering. This filtering step allows us to measure the constriction rate for each side independently. Constriction kinetics were derived by automatically extracting the fluorescent trajectories for ZipA and Pal using *KymographDirect*^93^. Anisotropy of the division process was determined by taking the ratio of the constriction velocity between the forward and reverse trajectory. Only cells where the division site displayed no signs of displacement except for constriction were analyzed to eliminate confounding effects on the analysis by excessive cell movement (e.g., pushing).

### Measuring division site circularity of vertically imaged cells

For vertical imaging of bacterial cells undergoing division, similar procedures as described previously^94,95^ were applied. Briefly, a silicon wafer containing 5.5 μm long and 1.5 μm wide photo-resist pillars was generated following high aspect ratio photolithography procedures with an adhesion layer. The dimension of these pillars reaches the practically feasible aspect ratio for photolithography designs and thus impedes increasing pillar length without concomitantly increasing width, precluding elongated or chaining division mutants. A modified silanization surface treatment with plasma cleaning was applied to increase the surface hydrophobicity of the silicon wafer to minimize agarose accumulation. Agarose micro holes were generated by pouring degassed 6 % agarose (w/v) in H_2_O on the silicon wafer. Agarose was allowed to solidify for 40 min at RT and was peeled of, cut into 5 × 5 mm pieces, and incubated in M9 medium supplemented with 0.2 % D-glucose, casamino acids, 25 μg ml^-1^ chloramphenicol and 50 μg ml^-1^ ampicillin overnight.

Cells were grown as described for sample preparation for live-cell imaging added on agarose pads. Cells which were not trapped in micro holes were washed off gently using 1 ml of growth medium. Five micrometer spanning Z-stacks (at a 200 nm step size) were acquired and subsequently deconvolved.

Circularity quantification was carried using the software package *Morphometrics*^96^. Fluorescence signals were segmented using Laplacian algorithm in combination the peripheral fluorescence setting. Circularity (*C*) is calculated in *Morphometrics* as:

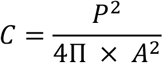

where (*P*) is the perimeter and (*A*) is the area enclosed by the circle and is a dimensionless measure. A perfect circle displays a circularity of 1, while increasing values correspond to less circular objects.

### Measuring Z-ring condensation from time-lapse data

Condensation of cytoskeletal elements was addressed using similar procedures described by Squyres et al., 2021. Briefly, five frames (corresponding to 10 min) from recorded time-lapse series were sum-projected in Fiji. Z-rings in these sum projection images were then aligned along the length axis and average-intensity projected into single image. Fluorescence intensity was measured across the full width along horizontal axis of the averaged projection image. Intensity values were normalized and their corresponding FWHM values were calculated in MATLAB.

### Measuring Z-ring condensation from 3D data

Similar procedures as outline for measuring Z-ring condensation in time-lapse series were applied. Two micrometer spanning Z-stacks (at a 200 nm step size) were acquired to capture a full three-dimensional view of a cell. Images were restored in Huygens as described above. Image volumes were sum-projected into a single plane, Z-rings extracted, aligned, and averaged described above. Fluorescence intensity profiles were measured identically as for time-lapse data. Snapshots for three-dimensional maximum intensity projections were rendered in Huygens.

### Measuring cell wall synthesis rates by biorthogonal NAM probes

Septal cell wall synthesis rates were measured as described previously here^98,99^. NAM-Alkyne was purchased as a custom synthesis product from Tocris following the procedures of Liang et al., 2017. All experiments were carried out in *ΔmurQ* background and in presence of pCF436^100^ for ITPG inducible expression of AmgK and MurU. Overnight cultures were back diluted 1:1000 into fresh LB containing 15 μg ml^-1^ gentamycin. Cells were grown at 37°C until OD_600_ = 0.4. Subsequently, 1.5 ml of cells were harvested (2 min, 5000 x g, RT) and resuspended in 300 μl LB containing 1 mM IPTG and 0.5 mM HADA to label all cell wall material by FDAAs. Samples were incubated rotating at 37°C for 30 min. Endogenous UDP-NAM production was inhibited by the addition of 200 μg ml^-1^ fosfomycin. After 10 min incubation, cells were washed twice in 1.5 ml LB, 1 mM IPTG, 200 μg ml^-1^ fosfomycin. Next, cells were incubated for 15 min in the presence of 0.2 % (w/v) NAM-Alkyne, 1 mM IPTG, 200 μg ml^-1^ fosfomycin at 37°C. Cells were fixed using ice-cold 70 % (w/v) ethanol for 20 min at 4°C. Next, cells pellets were washed 3x with 1x PBS. Biorthogonal NAM-Alkyne probes were labeled by click chemistry using 5 μM of Alexa488 azide substrate according to manufactures instruction. Samples were stored in 20 μl PBS at 4°C and imaged within 48 h of labeling experiment.

Samples were imaged on Nikon Ti2-E inverted widefield microscope equipped with a Lumencor Spectra III light engine, and Semrock dichroics (LED-CFP/YFP/mCherry-3X-A-000, LED-DA/FI/TR/Cy5/Cy7-5X-A-000) and emission filters (FF01-432/36, FF01-515/30, FF01-544/24). Images were recorded using a 1.45 NA Plan Apo 100 x PH3 oil objective with Olympus Type F immersion oil and a pco.edge 4.2bi Back Illuminated Cooled sCMOS camera using Nikon Elements 5.2.

One micrometer spanning Z-stacks (separated by 200 nm) were acquired and subsequently deconvolved as described under image processing for fluorescence microscopy. Z-stacks were sum-projected using Fiji. *De novo* septal PG synthesis was measured by integrating the fluorescence intensity of NAM-Alexa488 along the division site using the line tool (width = 3 pixels). Levels of cell wall hydrolysis were assessed by measuring the overall reduction in HADA fluorescence as compared to baseline signal intensity derived from fixing cells prior to NAM chase.

### Measuring cell wall remodeling by FDAA incorporation

For FDAA pulse chase experiments, cells grown overnight were back diluted 1:1000 in fresh LB and grown until OD600 = 0.4 at 37°C. Subsequently, 1.5 ml of cells were harvested (2 min, 5000 x g, RT) and resuspended in 300 μl LB containing 0.5 mM YADA. Samples were incubated rotating at 37°C for 40 min. Cells were washed once in 1.5 ml LB and resuspended in 300 μl LB containing 0.5 mM HADA. Samples were incubated at 37°C for either for 2 min, 4 min, and 8 min prior to immediate fixation with 70 % ethanol. After fixation, cells were washed 3x in PBS, stored in the dark at 4°C and imaged within 48 h. The same image acquisition and analyses procedures were carried out as described for NAM probes. In addition to the division site, fluorescence intensities measurements were also performed along the side wall and polar region of the cells.

### Cell shape quantification analyses

Bacterial cells were segmented and analyzed from still phase contrast images or time-lapse series using the software package *Morphometrics*^96^. Results from *Morphometrics* were post-processed using customized MATLAB scripts to exclude erroneously segmented cell debris in live image data based on area. Cell width, length and pole angles per segmented cell were directly extracted from *Morphometrics*. We obtained the invagination angle from both sides of the cell at the invagination site. The invagination site is defined as the narrowest segment of the cell, e.g., lowest cell width value, that present negative curvature on both sides of the cell body. The cell elongation (*E*) was determined by a customized MATLAB script:

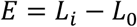

where *L* is cell length, *L*_0_ is the cell length in the first frame and i = 1, …, *I*, where *I* is the number of frames. The elongation rate is the slope of the linear regression performed on the plotted elongation values over time in Prism (GraphPad 9.0.0).

### SIM-TIRF Microscopy and MreB Tracking

Samples were prepared as described for live-cell imaging. Cells were added to high precision # 1.5 coverslips (Marienfeld) and placed on a 1 % (w/v) agarose pad in M9 (0.2 % D-glucose, casamino acids) and imaged at room temperature on a Nikon Ti-2 N-SIM microscope, equipped with N-SIM spatial light modulator illuminator, TIRF Lun-F laser combiner with 488 and 561 nm laser lines, a N-SIM 488/561 dual band dichroic mirror, SR HP Apo TIRF 100x 1.5 NA oil objective with automated correction collar and Hamamatsu Orca Flash 4.0 camera attached to a Cairn Research Twimcam splitter with a ET525/50m or a ET605/70m emission filter (for MreB-sw-mNeonGreen or Pal-mCherry fusion, respectively). The refractive index of the immersion oil (1.512) (GE Heatlhcare) was optimized for MreB-sw-mNeonGreen signal and corrected using the automated correction collar for the Pal-mCherry fusion. Alignment of the 488 and 561 lasers for SIM-TIRF and 3D-SIM and of the N-SIM optics and illumination was performed before each experiment at the image plane. First, a 3 min time-lapse series (at 3 s acquisition frame rate) in SIM-TRIF mode was collected using 20 % laser power with 100 ms exposure time to follow MreB-sw-mNeonGreen dynamics. Then, a single slice of a 3D-SIM Pal-mCherry (40 % laser power, 100 ms exposure) and a brightfield reference image was acquired. Raw fluorescence images were reconstructed using Nikon Elements 5.11 acquisition Software with indicated settings: MreB illumination contrast 0.8, noise suppression 0.3 and blur suppression 0.05; Pal illumination contrast 3.75, noise suppression 0.1 and blur suppression 0.5. Only reconstructed images with a quality score ≥ 8 and passed SIMcheck quality test^101^ were used for further analysis. Subsequently MreB time-lapse series was overlayed over the reference channels in Fiji.

Particle tracking was performed in Fiji using the TrackMate v6.0.1 plugin^102^. MreB filaments were detected using the LoG-detector with an estimated radius of 0.3 μm. Spurious spots were filtered using a quality threshold of 50. Spots were linked using a Kalman filter with an initial search radius for 0.2 μm and search radius for 0.1 μm. No frame gaps were allowed. Only Tracks consisting of ≥ 4 continuous spots (12 s) and traveled less than 1 μm in total distance were kept for further analysis. To analyze the nature of the displacement of each track, the mean square displacement (*MSD*) was calculated using the MATLAB class msdanalyzer^103^ following the equation:

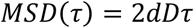

where (*t*) is the delay time and (*D*) is the diffusion coefficient. Slopes (*α*) of the individual *MSD* curves were extracted using the Log-log fit of the *MSD* and the delay time *τ*. As the maximum delay time 75 % of the track length was used. Tracks with a *R*^2^ for log [*MSD*] versus log [*t*] below 0.95 indicative of a poor fit to the MSD curve were excluded from the analysis. MreB filaments engaged in active cell wall synthesis are displaced by the enzymatic action of the enzymatic activities of RodA and PBP2b^56–58^ and thus it’s *MSD* curves display slopes of *α* ≈ 2 indicative of a transported particle motion above the rate of Brownian diffusion (Fig. S10B). MreB filaments in constricting cells, as determined by the presence of a Pal-mCherry foci at the division site, were analyzed by fitting a 200 nm wide region of interest to the cell division site. Directional MreB tracks were deemed to contribute to the elongation of the division site. Early and late division stages were distinguished by the presence of two separated Pal foci or a continuous fluorescent signal across the cell, respectively.

### Statistical analysis

All data measurements were plotted and analyzed using GraphPad Prism 9 (Version 9.1.2). In general, (log-) normal distribution was tested by using Shapiro-Wilk test, for comparisons of two groups, significance was determined by two-tailed, unpaired Student’s t test with Welch correction and F test for variance analysis. One-way ANOVA test were used for comparison of more than two groups using the recommended post-test for selected pairwise comparisons. All experiments were carried out at least with 3 independent biological replicates. P values less than 0.05 were considered statistically significant. Levels of significance are indicated by *p<0.05, **p<0.01, ***p<0.001, ****p<0.0001; ns, not significant.

