## Supplementary Material for "Cell wall synthesis and remodeling dynamics determine bacterial division site architecture and cell shape"

Thomas G. Bernhardt

Department of Microbiology, Harvard Medical School, Boston, USA

Luke H. Chao

Department of Molecular Biology, Massachusetts General Hospital, Boston, USA

Department of Genetics, Harvard Medical School, Boston, USA

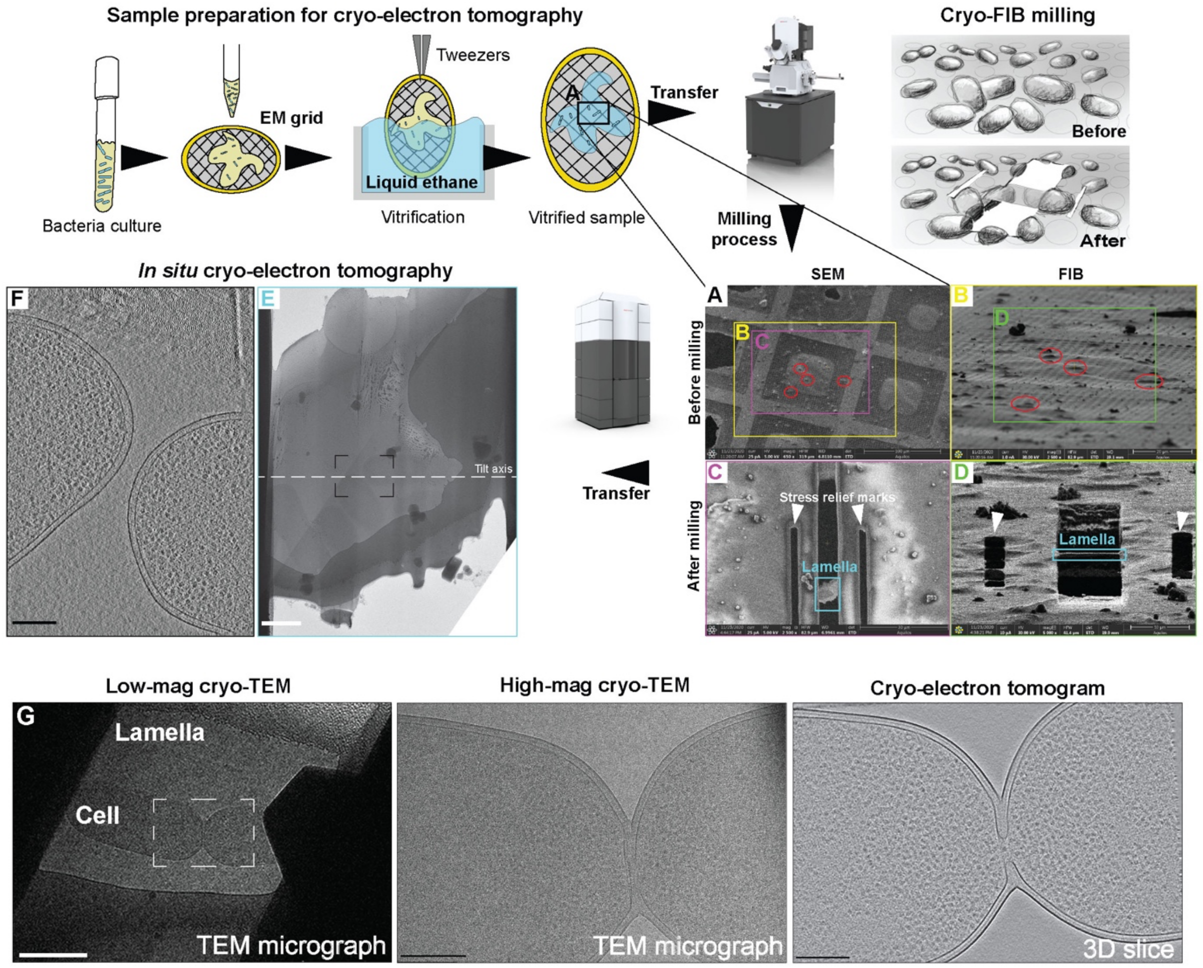


**Supplementary Figure 1: Cryo-FIB / cryo-ET pipeline utilized in this study**. Schematic cartoons showing the steps in sample preparation for cryo-ET. In brief, bacteria are grown to $OD_{600}$ = 0.3 and applied onto an EM grid for vitrification in liquid ethane (Dubochet et al., 1988). Cryo-EM grids are kept in liquid nitrogen until transfer into the cryo-FIB microscope for milling. An illustration shows the result of milling vitrified bacteria distributed onto the holey carbon film on the mesh EM grid. (A-D) Images taken from the Aquilos Thermo Fisher Scientific graphical user interface during cryo-FIB milling performance. (A) Target bacteria (red circles) are first identified by SEM, $e^-$ beam. Yellow box indicates region visualized in (B) and magenta box indicates the area visualized in (C). (B) Corresponding FIB, ion $Ga^+$ beam, view (52° with respect to the $e^-$ beam (Wagner et al., 2020)) of the targeted grid square in (A) (yellow box). Green box indicates region visualized in (D). (C) SEM view of the same region shown in (A) and (B) after platinum deposition and milling. Cyan box indicates obtained lamella shown in (E). (D) FIB view of region shown in (B) (green box) after platinum deposition and milling. Scale bars in (A-D) are indicated on each image. After milling, cryo-EM grids containing bacterial lamellae are transferred into a TEM microscope. (E) Low magnification TEM 2D image of the lamella shown in (C-D), cyan box. Dashed black box indicates target region for cryo-ET acquisition. Dashed white line indicates the tilt axis for cryo-ET data acquisition. (F) 3D slice of the cryo-electron tomogram obtained from 3D reconstruction of aligned cryo-ET tilt series acquired in (E) (dashed black box). Scale bars: E = 1000 nm; F = 200 nm. (G) A representative lamella from a wild-type *E. coli* cell imaged at indicated imaging conditions. White box highlights region for corresponding high-magnification acquisition. Scale bars = 1 µm (low magnification); 200 nm (high magnification and cryo-electron tomogram).

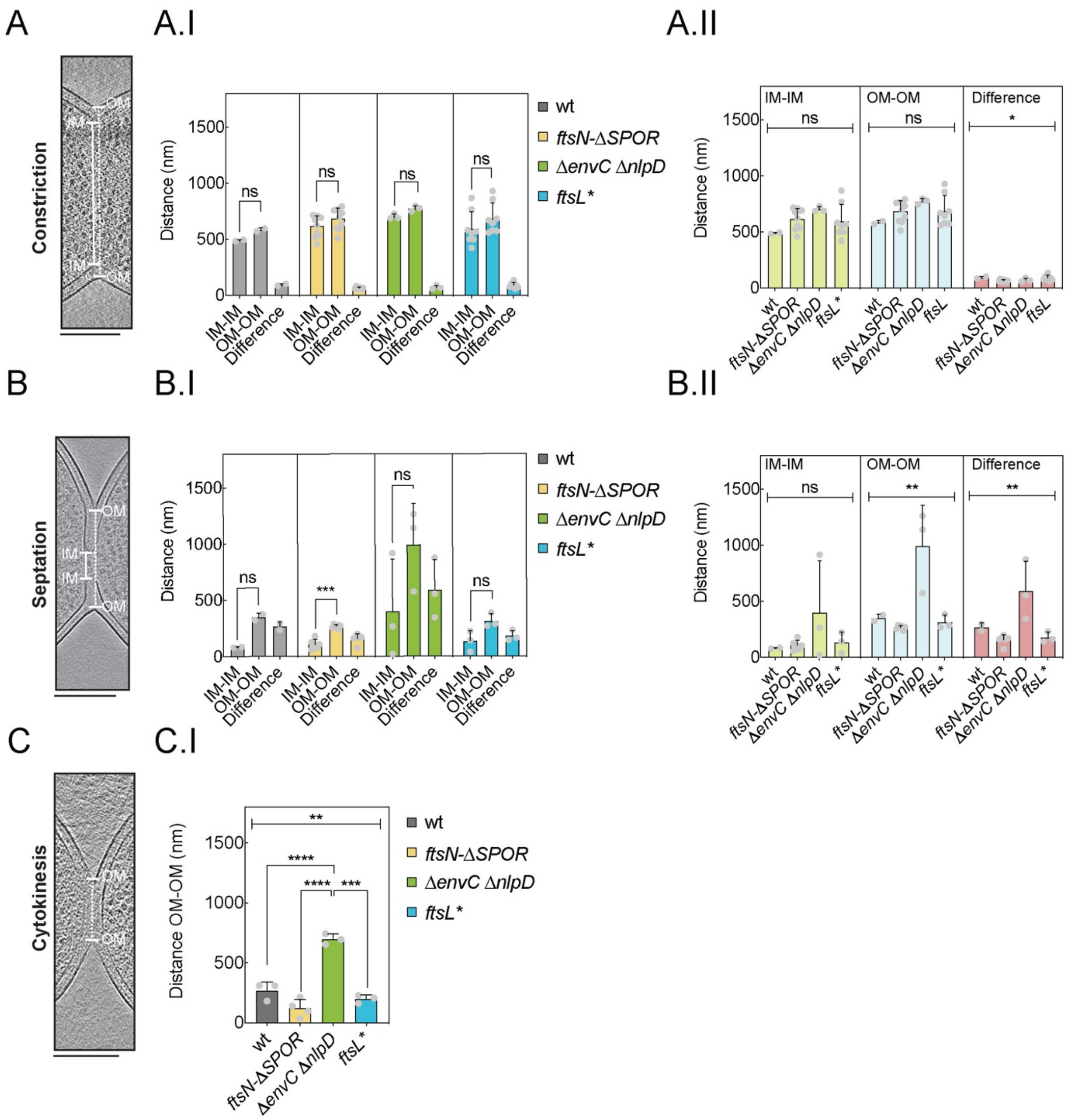


**Supplementary Figure 2: Quantification of division site distances in cryo-ET data of dividing *E. coli*.** 3D slices visualizing the division site during (A) constriction, (B) septation and (C) cytokinesis. Dashed white line indicates OM-OM distance and white bold line indicates IM-IM distance. (A.I-C.I) show bar graphs plotting the measured distances in nm of IM-IM, OM-OM and the difference between OM-OM and IM-IM distances at (A.I) constriction, (B.I) septation and (C.I) cytokinesis stages for each strain. (A.II-B.II) Bar graphs plotting distances in nm for each strain grouped as IM-IM distance, OM-OM distance and the difference of OM-OM and IM-IM distance at (A.II) initiation and (B.II) active constriction stages. Scale bars = 200 nm. All data are expressed as mean ± SEM. * = $p < 0.05$, ** = $p < 0.01$, *** = $p < 0.001$, **** = $p < 0.0001$.

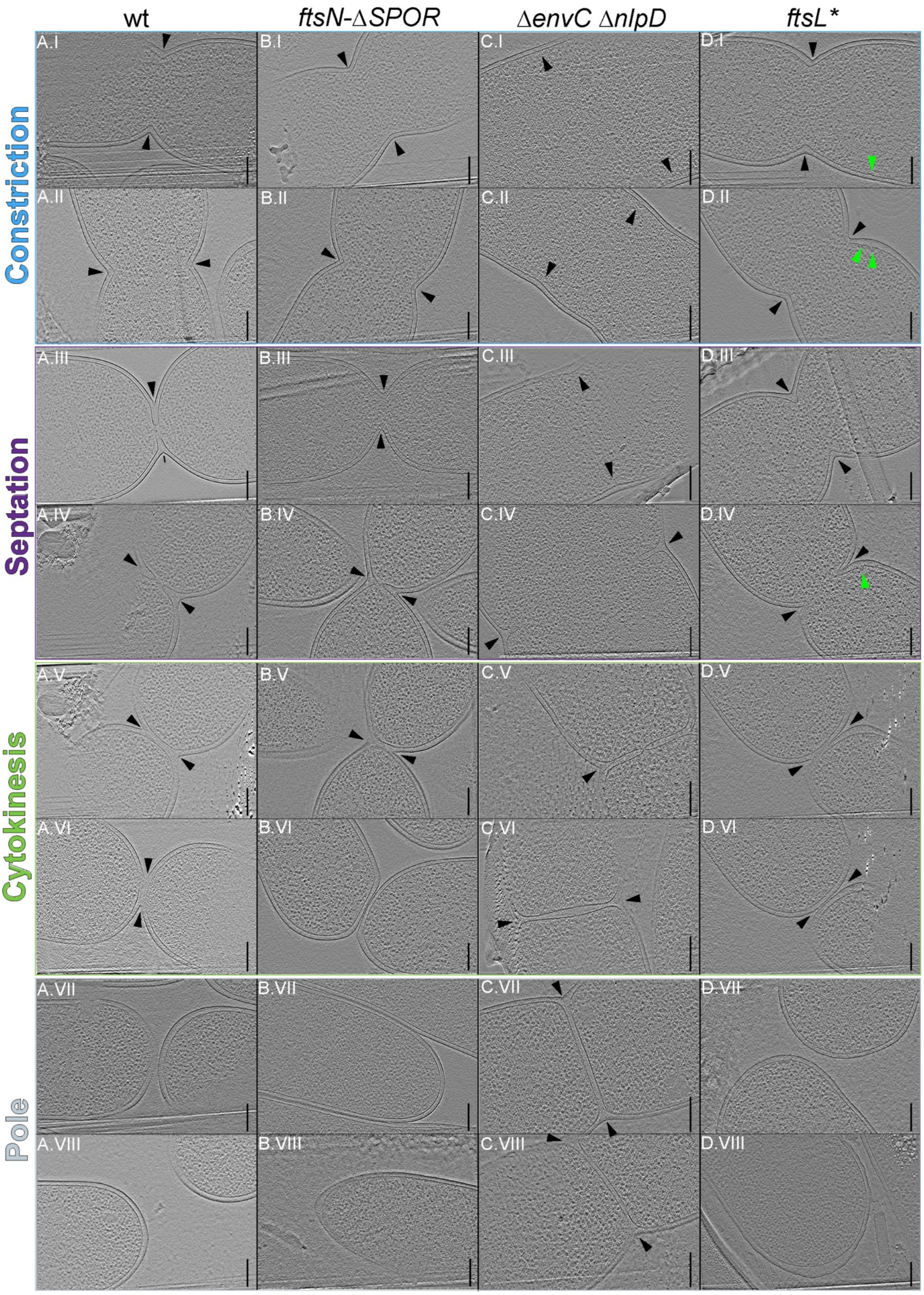


**Supplementary Figure 3: Cell division and polar morphology of *E. coli* viewed by cryo-ET.** Gallery of summed projected central slices of cryo-electron tomograms visualizing the indicated division mutants. Black arrowhead = division site; green arrowhead = IM bulging. Scale bars = 200 nm.

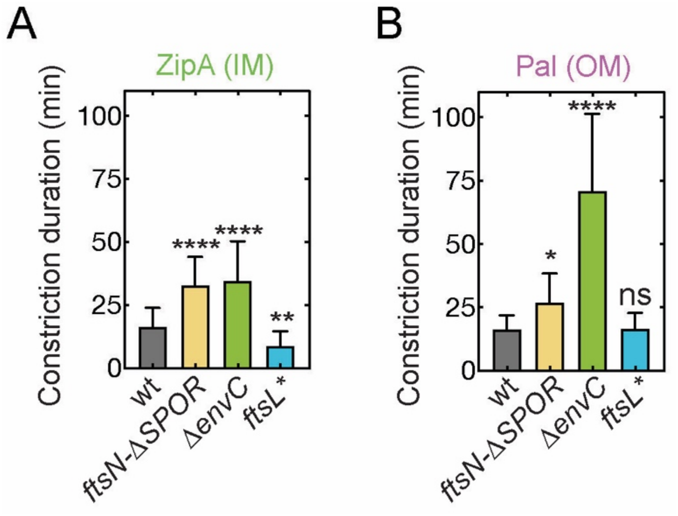


**Supplementary Figure 4: Duration of cell envelope constriction during cell division.** Duration of (A) IM and (B) OM constriction was derived from kymograph measurements as stated in Fig. 2C-D. Data are represented as mean + SD. One-way ANOVA, differences in significances are tested relative to wild-type; ns = non-significant, * = $p < 0.05$, *** = $p < 0.001$, **** = $p < 0.0001$; N = 150.

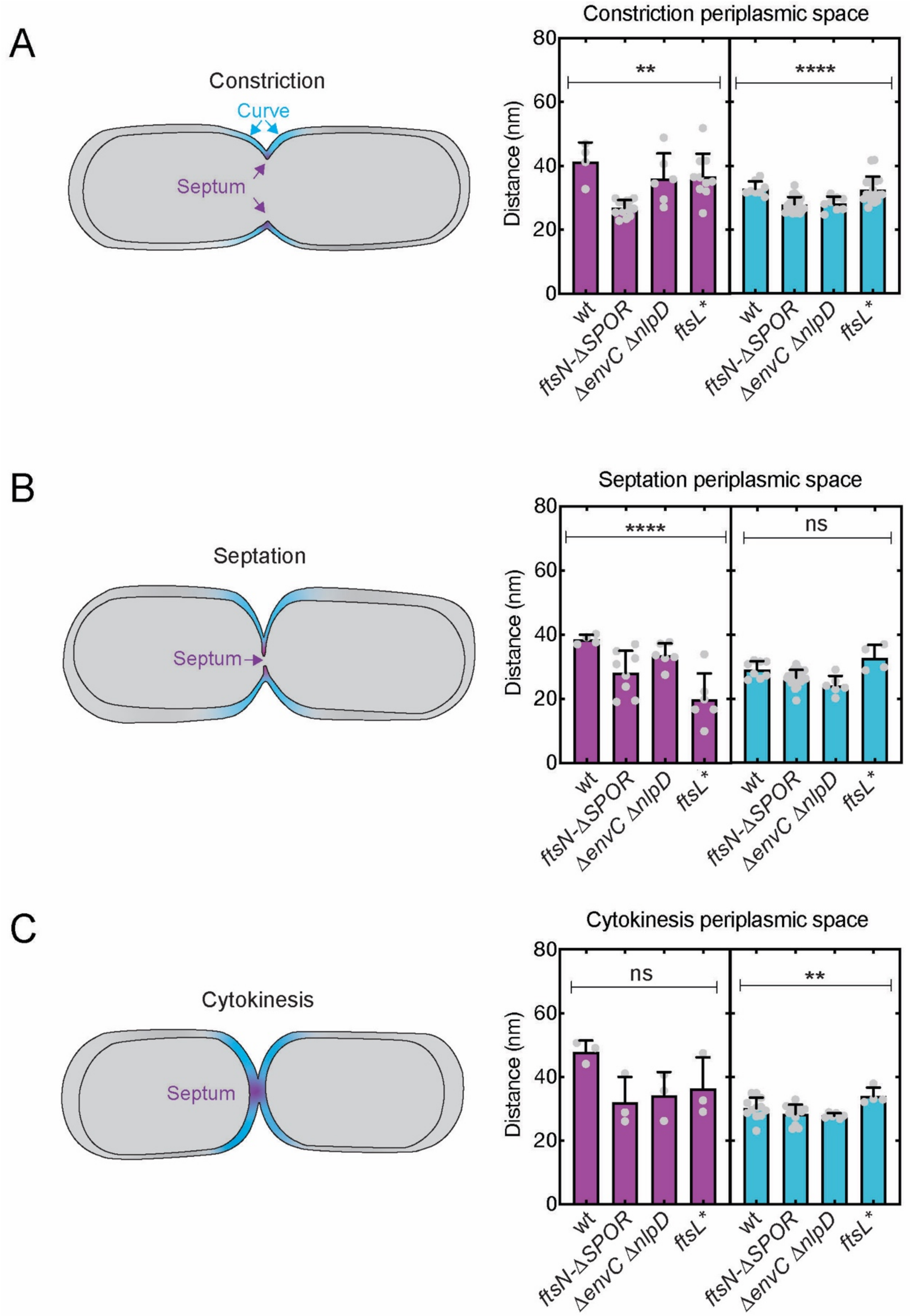


**Supplementary Figure 5: Quantification periplasmic width in dividing *E. coli* from cryo-ET data.** (A-C) Schematic cartoon representing the division stages of *E. coli* and indicating the color-coded regions where periplasmic space width was measured. Corresponding bar graph of periplasmic space width measured in the specified color-coded regions shown in a of the indicated division mutants. All analyzed data points are displayed, bar represents mean ± SEM. Brown-Forsythe and Welch ANOVA test, significance was tested among all groups within each region; ns = non-significant, ** = $p < 0.01$, **** = $p < 0.0001$.

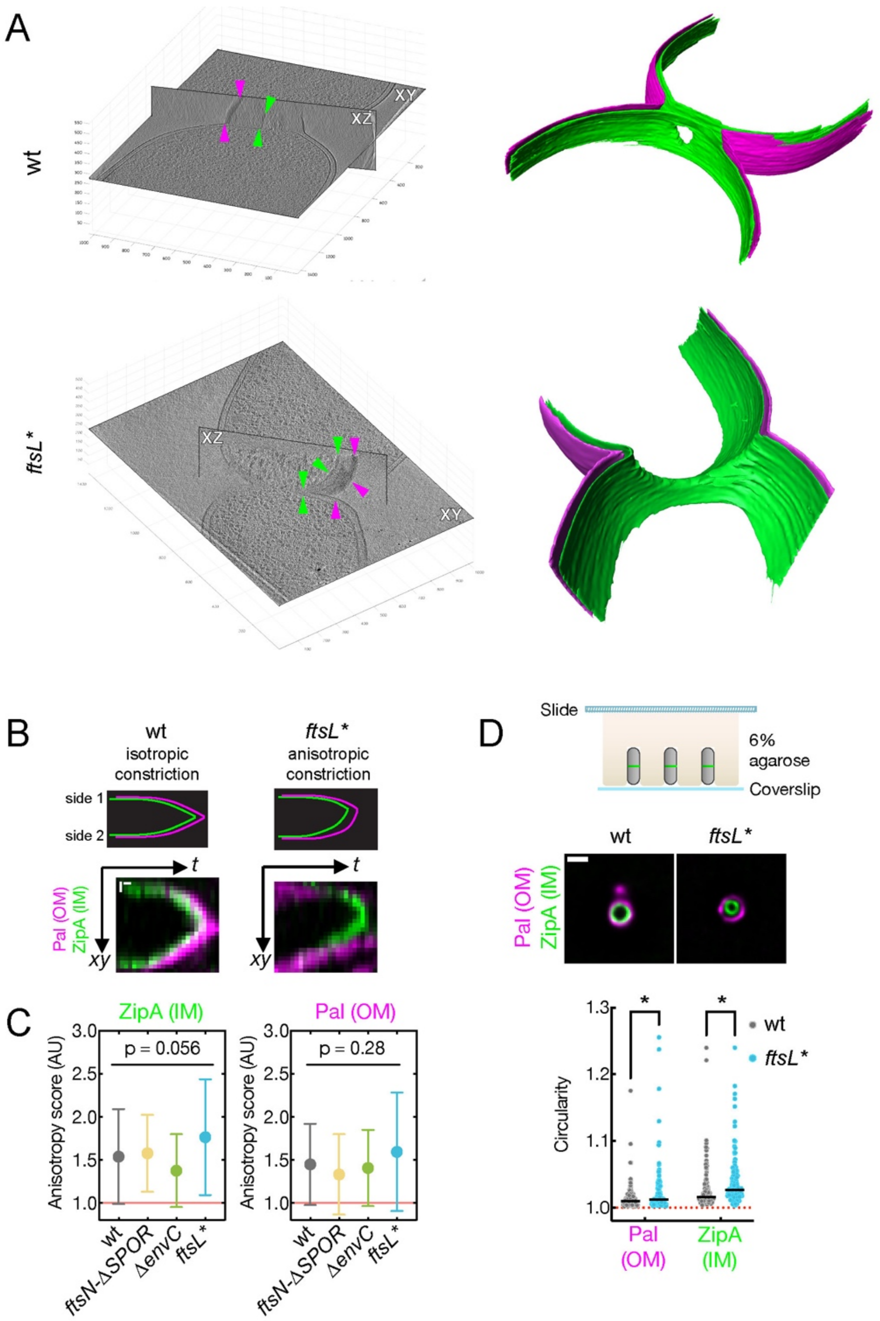


**Supplementary Figure 6: A hyperactivated divisome leads to anisotropic cell envelope constriction.** (A) Orthogonal views of XZ and XY slices of 3D cryo-electron tomograms of the indicated division mutants. Magenta and green arrowheads indicate OM and IM, respectively. 3D volumes are displayed in cartesian 3D grids with axes indicating the dimensions in pixels. For WT 100 pixels = 102.6 nm, and for *ftsL** 100 pixels = 110.3 nm. Corresponding 3D surface segmentation renderings of OM (magenta) and IM (green) are shown on the right. (B) Schematic overview of a theoretical kymograph for an isotropic (left) and anisotropic (right) constriction of the cell envelope. Real examples for wild-type (left) and *ftsL** (right) are provided. (C) An anisotropy score was calculated by taking the ratio of the constriction velocity from both sides of the cell. Red line (= 1) indicates a perfectly isotropic cell envelope constriction process. Data are represented as mean ± SD, Kruskal-Wallis test, p values are shown. (D) Cells were vertically immobilized using small micro pillars imprinted into agarose pads, allowing to image the cell division site along its long axis. Representative example of the cell envelope position in vertically imaged wt and *ftsL** cells. Scale bar = 2 μm. Circularity was quantified using *Morphometrics*. Red line (circularity = 1) indicates a perfect circle. Two-way ANOVA with Sidak's multiple comparison test, * = $p < 0.05$.

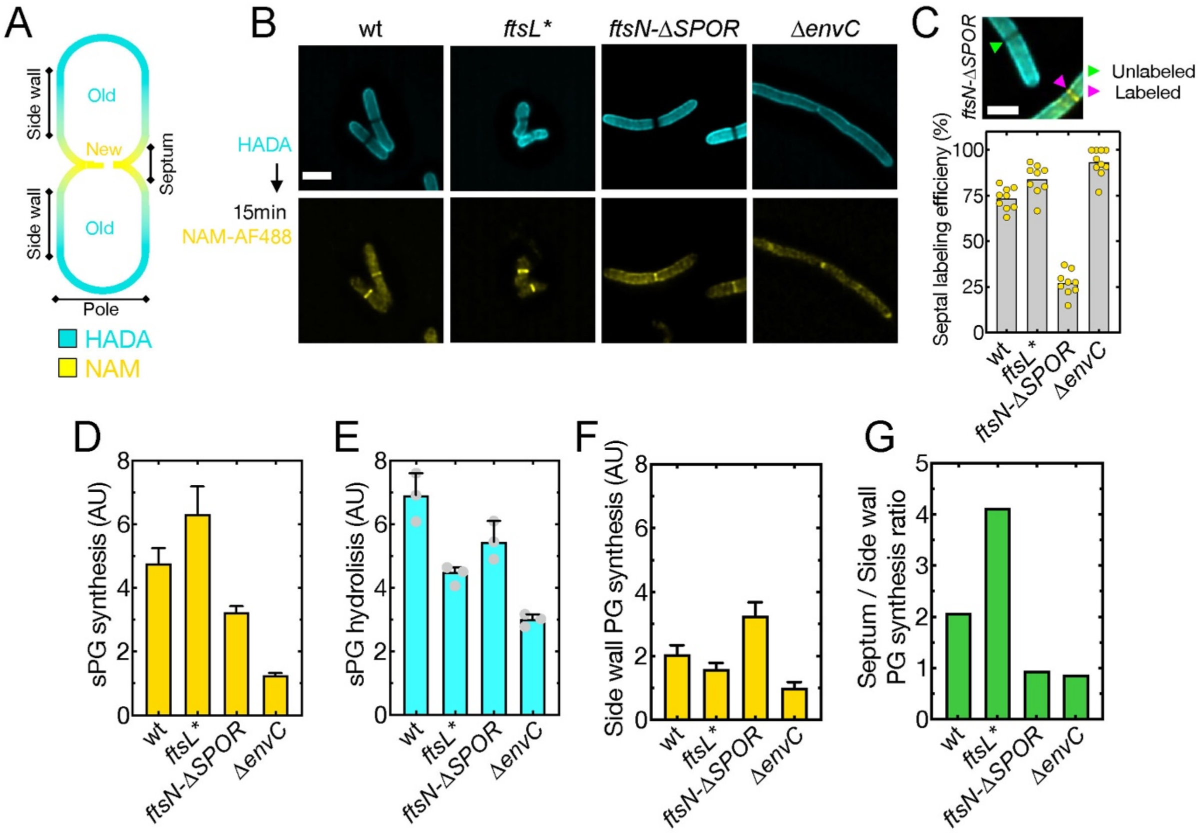


**Supplementary Figure 7: Measuring cell wall synthesis and hydrolysis during division and elongation using a biorthogonal NAM probe**. (A) Schematic representation of labelling patterns for the NAM-FDAA based pulse-chase experiment. New cell wall material will be labeled by Alexa488 labeled NAM (yellow), while old material is stained by HADA (blue). (B) Sum-projected deconvolved images of indicated strains labeled cells after a 15 min pulse-chase with NAM. (C) Bar graph assessing NAM incorporation at the division site. Example for a labeled and unlabed division site in *ftsN*-Δ*SPOR* mutant is shown. (D) Rate of sPG synthesis was determined by measuring integrated fluorescence intensities of NAM and normalized to division site area. (E) Rates of sPG hydrolysis were determined by calculating the average reduction in HADA fluorescence compared to cells fixed prior to the 15min NAM chase. Points indicate the average of three biological replicates. (F) Side wall NAM incorporation was determined by measuring integrated fluorescence intensities of NAM and normalized to area. Septal to side wall PG synthesis ratios were calculated by dividing the average fluorescence intensity values for septal synthesis by values for sidewall synthesis. All data are represented as mean with 95% CI shown as error bars. All values are in arbitrary units and were divided by 1000 for plotting purposes. Scale bars = 2μm.

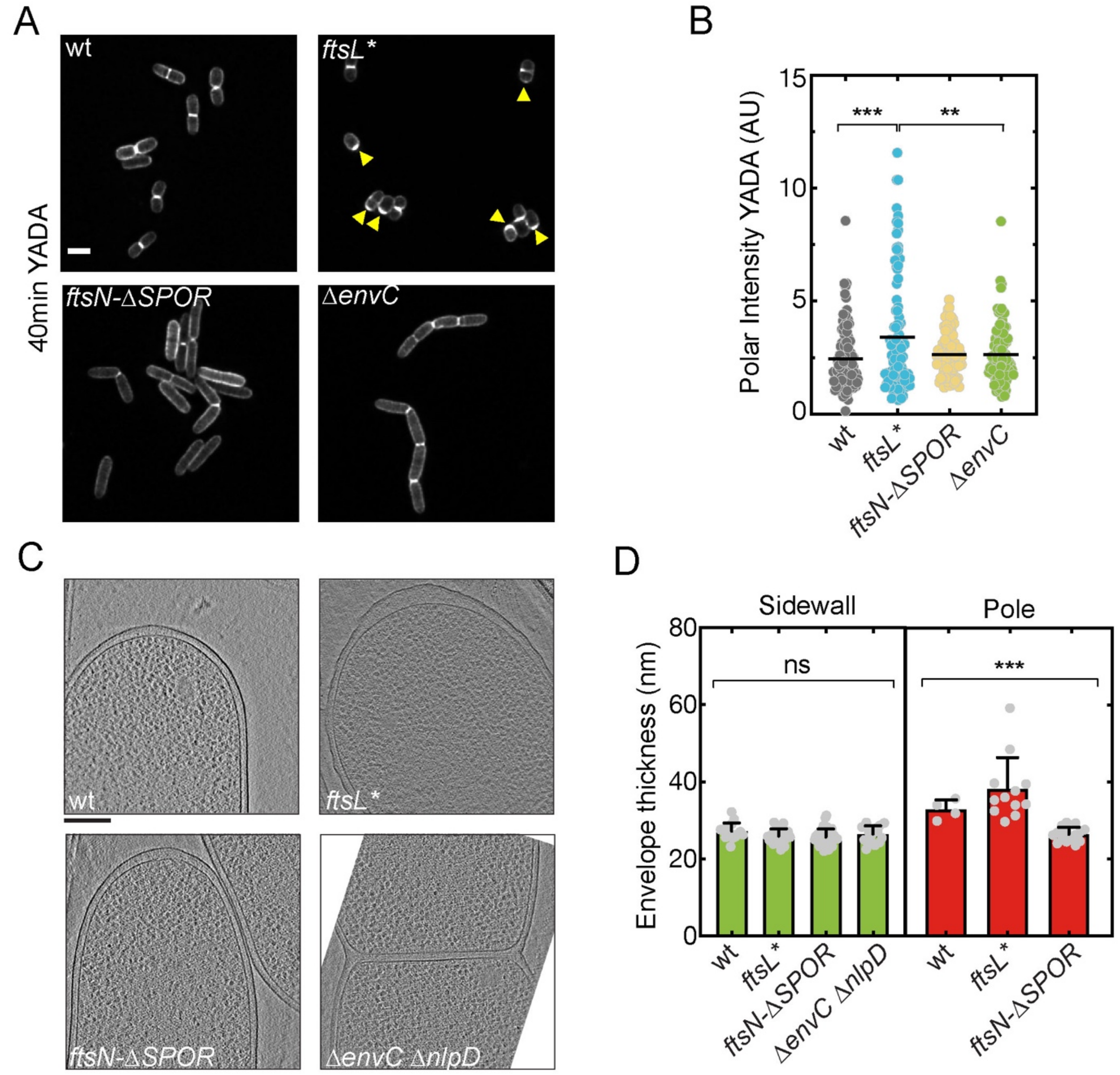


**Supplementary Figure S8: The *ftsL** mutant displays elevated intensities of cell wall material and an enlarged periplasmic space at the cell pole.** (A) Sum projection of a 1 µm spanning z-stack of YADA labeled division mutants. Yellow arrow heads indicate accumulation YADA at cell poles in cells, which presumably recently separated. Scale bar = 2 µm. (B) Polar YADA fluorescence intensity was integrated normalized by area. Values are in arbitrary units and were divided by 1000 for plotting purposes. Individual data points and mean are displayed. One-way ANOVA, ** = $p < 0.01$, *** = $p < 0.001$, N = 100 cells. (C) Summed projected central 3D slices through cryo-electron tomograms containing cell poles. All images are scaled; scale bar = 200 nm**.** (D) Bar graph showing the measured periplasmic space thickness in cryo-ET data at the cell side wall (green) and pole (red) for each indicated mutant. All data are expressed as mean ± SEM. One-way ANOVA; significance was tested among all groups within each region; ns = non-significant, *** = $p < 0.001$.

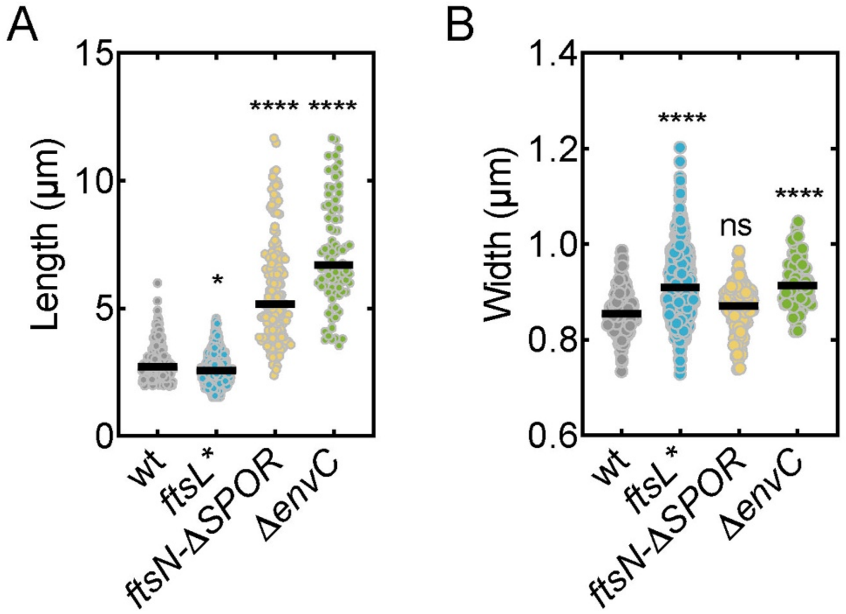


**Supplementary Figure 9: Cell length and width measurements.** Median cell length (A) and width (B) was measured for the indicated divisome mutants using Morphometics. Kruskal-Wallis One-way ANOVA, differences in significances are tested relative to wild-type, * = p < 0.05, **** = p < 0.0001, ns = non-significant, N = 230 cells.

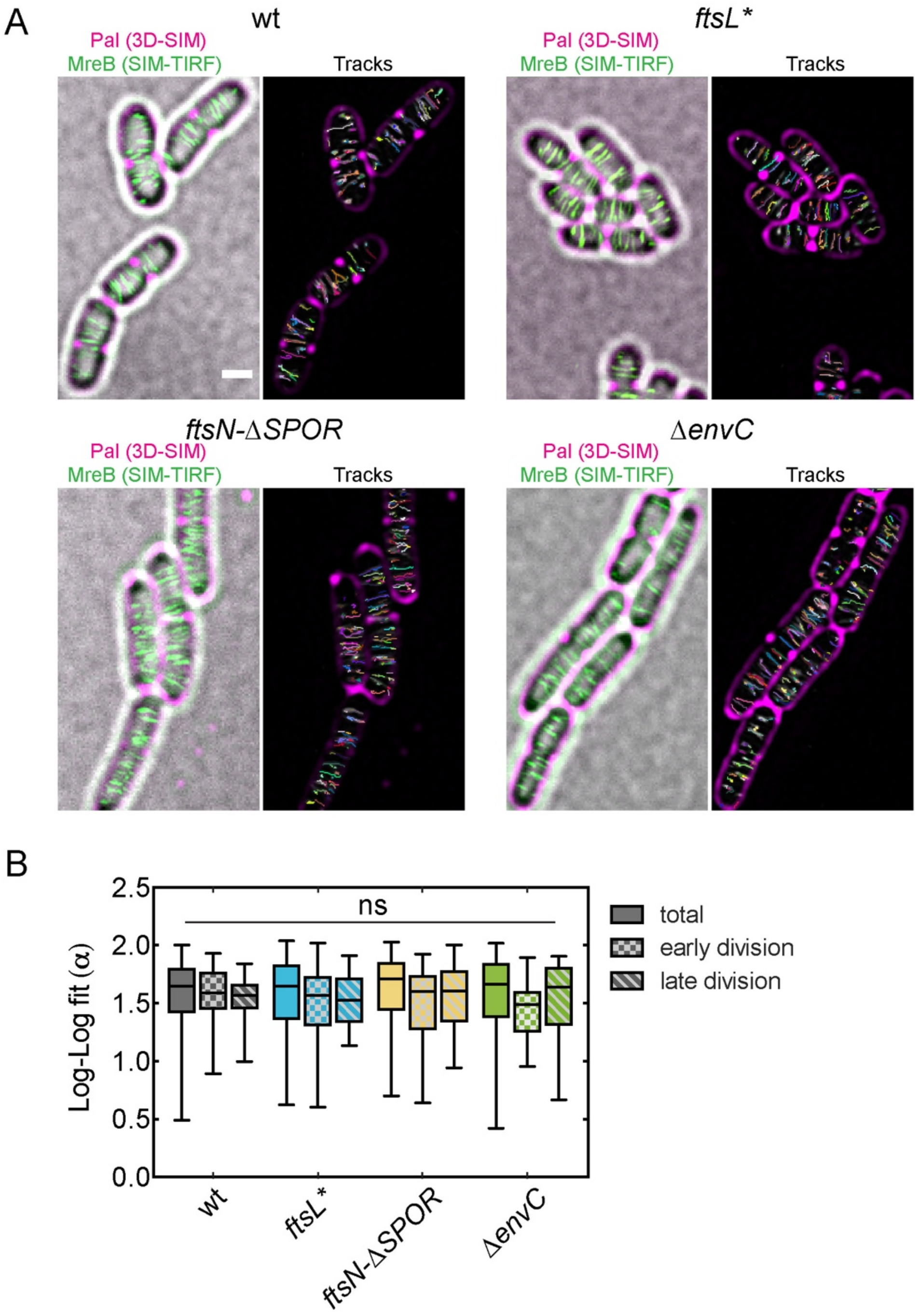


**Supplementary Figure 10: MreB tracking shows Rod complex activity at the division site in constricting cells.** (A) MreB-sw-mNeonGreen dynamics were followed by SIM-TRIF microscopy for 3 min at 3 s acquisitions per frame in indicated mutants. Time-lapse series was sum-projected and overlayed over a 3D-SIM Pal-mCherry and brightfield reference image. Tracking results from TrackMate are displayed on the right. For clarity, display of brightfield image was omitted and MreB filaments are shown in grayscale. Cells were imaged at room temperature on M9 supplemented with 0.2 % glucose and casamino acids. Larger fields of view are shown as compared to Figure 5G. (B) Slopes of MSD curves $(\alpha)$ were analyzed following log-log fit to $\log[MSD]$ versus $\log[t]$ using the MATLAB class msdanalyzer. Particles displaced by diffusive motion are characterized by a slope of their $\log[MSD]$ = 1, while transported particles have slopes of 2 and constrained particles display slopes < 1. Box plot error bars displaying Min-Max range of values. Fifty percent of values lie within the box, blackline represents median. Kruskal-Wallis test, N = 2588 tracks were analyzed, significance was determined against wild-type; ns = non-significant. Scale bar = 1 µm.

**Supplementary Table 1. Summary of data acquisition and image processing for cryo-ET data in this study.**

| Sample | | wt | *ftsL** | *ftsN-ΔSPOR* | *ΔenvC ΔnlpD* |
|---|---|---|---|---|---|
| **Cryo-FIB milling** | **Microscope** | Aquilos Cryo-FIB, FEI – Thermo Fisher Scientific | Aquilos Cryo-FIB, FEI – Thermo Fisher Scientific | Aquilos Cryo-FIB, FEI – Thermo Fisher Scientific | Aquilos Cryo-FIB, FEI – Thermo Fisher Scientific |
| **Acquisition settings** | **Microscope** | Titan Krios Gi3 FEI, Thermo Fisher Scientific | Titan Krios Gi3 FEI, Thermo Fisher Scientific | Titan Krios Gi3 FEI, Thermo Fisher Scientific | Titan Krios Gi3 FEI, Thermo Fisher Scientific |
| | **Voltage (KeV)** | 300 | 300 | 300 | 300 |
| | **Detector** | Gatan K3 IS | Gatan K3 IS | Gatan K3 IS | Gatan K3 IS |
| | **Energy filter** | Gatan BioQuantum K3 | Gatan BioQuantum K3 | Gatan BioQuantum K3 | Gatan BioQuantum K3 |
| | **Slit width (eV)** | 20 | 20 | 20 | 20 |
| | **Super-resolution mode** | Yes | Yes | Yes | Yes |
| | **Å/pixel** | 1.282/1.379* | 1.379 | 1.379 | 1.282/1.379* and 1.096** |
| | **Defocus (µm)** | -3.5 to -5.0 | -3.5 to -5.0 | -3.5 to -5.0 | -3.5 to -5.0 |
| | **Acquisition scheme** | -70/70, 2°, Dose-symmetric | -70/70, 2°, Dose-symmetric | -70/70, 2°, Dose-symmetric | -70/70, 2°, Dose-symmetric |
| | **Total dose** | ~90 - 120 | ~90 - 180 | ~90 - 180 | ~90 - 120 |
| | **Dose rate (e-/Å/sec)** | ~ 1.5 - 3 | ~ 1.5 - 3 | ~ 1.5 - 3 | ~ 1.5 - 3 |
| | **Frame number** | 4 - 6 | 4 - 6 | 4 - 6 | 4 - 6 |
| | **Number of tomograms** | 9 | 16 | 27 | 8 |
| **Image processing** | **Frame alignment and dose weighting** | *framealign*, IMOD | *framealign*, IMOD | *framealign*, IMOD | *framealign,* IMOD |
| | **Tilt series alignment** | IMOD/*Dynamo* | IMOD | IMOD | IMOD |
| | **WBP** | IMOD | IMOD | IMOD | IMOD |
| | **Filtering** | IMOD/*Dynamo*/ Amira | IMOD/*Dynamo*/ Amira | IMOD/*Dynamo/* Amira | IMOD/*Dynamo*/ Amira |
| | **3D-segmentation** | Amira | Amira | Amira | Amira |
| | **3D-rendering** | *Dynamo*/Amira | *Dynamo*/Amira | *Dynamo*/Amira | *Dynamo*/Amira |

*Data was acquired in two different FEI Titan Krios (Thermo Fisher Scientific).
**Data was acquired at two different magnifications (35kx and 42kx).

**Supplementary Table 2. Strains used in this study.**

| Strain | Genotype[a] | Relevant features | Used for | Source/Reference[b] |
|---|---|---|---|---|
| TB28 | *rph1 ilvG rfb-50 ΔlacIZYA<>frt* | lacZ- MG1655, wild type | cryoET / FDAA | (Bernhardt and Boer, 2003) |
| TB44 | *TB28 ΔenvC<>frt* | *envC* deletion | | (Bernhardt and de Boer, 2004) |
| TB156 | *TB28 ΔenvC<>frt ΔnlpD::aph* | *nlpD* and *envC* double deletion, Kan$^R$ | | (Uehara et al., 2009) |
| TT154 | *TB28 ftsN(1-243)-TAA<>frt* | *ftsN-ΔSPOR* deletion | | (Truong et al., 2020) |
| MT10 | *TB28 ftsL(E88K)* | *ftsL** allele | | (Tsang and Bernhardt, 2015) |
| AV92 | *TB28 murQ::aph* | *murQ* deletion, pCF436 | NAM | P1(CF491)(Fumeaux and Bernhardt, 2017) x TB28 |
| AV93 | *TB28 ftsN(1-243)-TAA<>frt murQ::aph* | *ftsN-ΔSPOR* deletion in Δ*murQ*, pCF436 | | P1(CF491)(Fumeaux and Bernhardt, 2017) x TT154 |
| AV94 | *TB28 ΔenvC<>frt murQ::aph* | Δ*envC* deletion in Δ*murQ*, pCF436 | | P1(CF491)(Fumeaux and Bernhardt, 2017) x TB44 |
| AV125 | *TB28 ftsL(E88K) murQ::aph* | *ftsL** allele in Δ*murQ*, pCF436 | | P1(CF491)(Fumeaux and Bernhardt, 2017) x MT10 |
| AV53 | *TB28 attHK022* $P_{lac}$*::zipA-sfGFP bla pal-mCherry cat* | GFP-IM/mCherry-OM, Amp$^R$, Cam$^R$ | Live-cell imaging | P1(MG11) x TB28(attHKTB225)(Uehara et al., 2009) |
| AV67 | *AV53 ftsN(1-243)-ΔSPOR::aph* | *ftsN-ΔSPOR* deletion in GFP-IM/mCherry-OM, Amp$^R$, Cam$^R$, Kan$^R$ | | P1(NP279)(Truong et al., 2020) x AV53 |
| AV76 | *AV53 ΔenvC::aph* | *envC* deletion in GFP-IM/mCherry-OM, Amp$^R$, Cam$^R$, Kan$^R$ | | P1(JW5646)(Baba et al., 2006) x AV53 |
| AV81 | *TB28 attHK022 Plac::zipA-sfGFP ΔenvC::aph* | Strain-intermediate towards AV96 | | TB28(attHKTB225)(Uehara et al., 2009) x P1(JW5646)(Baba et al., 2006) |
| AV115 | *AV53 leu::TN10* | Strain intermediate for AV116, Amp$^R$, Cam$^R$, Tet$^R$ | | P1(CH43)(Liu et al., 2015) x AV53 |
| AV116 | *AV53 ftsL(E88K)* | *ftsL** allele in GFP-IM/mCherry-OM, Amp$^R$, Cam$^R$ | | P1(MT10)(Tsang and Bernhardt, 2015) x AV115 |

| | | | | |
|---|---|---|---|---|
| AV134 | *AV53 ftsL(E88K)* Δ*envC::aph* | Δ*envC* deletion in ftsL* in GFP-IM/mCherry-OM, Amp$^R$, Cam$^R$, Kan$^R$ | | P1(JW5646)(Baba et al., 2006) x AV116 |
| AV136 | *AV53* Δ*fliO::aph* | Flagellum deletion in GFP-IM/mCherry-OM, Amp$^R$, Cam$^R$, Kan$^R$ | | P1(JW5316)(Baba et al., 2006) x AV53 |
| AV137 | *AV53 ftsL(E88K)* Δ*fliO::aph* | Flagellum deletion in *ftsL** GFP-IM/mCherry-OM, Amp$^R$, Cam$^R$, Kan$^R$ | | P1(JW5316)(Baba et al., 2006) x AV116 |
| AV150 | *TB28 mreB'-mNeonGreen-'mreB ΔyhdE pal-mCh cat* | MreB-mNG, Pal-mCh, Cam$^R$ | SIM-TIRF | P1(MG11) x AV7 |
| AV151 | *AV150 ftsL(E88K)* | *ftsL** in MreB-mNG Pal-mCh background, Cam$^R$ | | P1(MG11) x AV149 |
| AV152 | *AV150* Δ*envC::aph* | Δ*envC* in MreB-mNG Pal-mCh background, Cam$^R$, Kan$^R$ | | P1(MG11) x AV146 |
| AV154 | *AV150 ftsN(1-243)-*Δ*SPOR::aph* | *ftsN-*Δ*SPOR* in MreB-mNG Pal-mCh background, Cam$^R$, Kan$^R$ | | P1(MG11) x AV147 |

[a] The KanR cassette is flanked by FLP recognition target (frt) sites for removal by FLP recombinase. An frt scar remains following removal of the cassette using FLP recombinase expressed from pCP20. Numbers in parentheses indicate the codons included in the relevant clones.

[b] Strain constructions by P1 transduction are described using the shorthand: P1(donor) x recipient. Transductants were selected on LB Kan, Tet, Cm, or minimal medium with no casamino acids plates where appropriate. Strains resulting from the removal of a drug resistance cassette using pCP20 (Datsenko and Wanner, 2000a) are indicated as: Parental strain/pCP20.

### Supplementary Table 3. Plasmid used in this study.

| Plasmid | Genotype[a] | ori | Relevant features | Source/Reference[b] |
|---|---|---|---|---|
| CF436 | *aacC1 bla Tn7 lacIq* $P_{lac}$*::amgK-murU* | colE1 | lacZ inducible AmgK and MurU for NAM | (Fumeaux and Bernhardt, 2017) |

**Supplementary Video 1. *In situ* cell division of wild-type *E. coli*.** Cryo-electron tomograms of wt *E. coli*. Time-lapse series were acquired with a rate of 7 fps in the compressed format MOV for visualization purposes. Green, cyan and magenta layers indicate segmented IM, PG and OM, respectively. Scale bars = 100 nm.

**Supplementary Video 2. Fluorescence live-cell imaging of cell envelope constriction in *E. coli* division mutants.** Three time-lapse series of each indicated *E. coli* mutants acquired at a 2:30 min:sec acquisition interval are shown. Bacteria were imaged at 30˚C on 1 % agarose in M9 supplemented with 0.2 % casamino acids and D-glucose. Fluorescence channels (Pal-mCherry – magenta, ZipA-sfGFP – green) were deconvolved. Video was rendered at 12 fps. Scale bar = 2 µm.

**Supplementary Video 3. *In situ* cell division of *ftsN-ΔSPOR*.** Cryo-electron tomograms of *ftsN-ΔSPOR* mutant. Time-lapse series were acquired with a rate of 7 fps in the compressed format MOV for visualization purposes. Green, cyan and magenta layers indicate segmented IM, PG and OM, respectively. Scale bars = 100 nm.

**Supplementary Video 4. *In situ* cell division of *ΔenvC ΔnlpD*.** Cryo-electron tomograms of *ΔenvC ΔnlpD* mutant. Time-lapse series were acquired with a rate of 7 fps in the compressed format MOV for visualization purposes. Green, cyan and magenta layers indicate segmented IM, PG and OM, respectively. Scale bars = 100 nm.

**Supplementary Video 5. *In situ* cell division of *ftsL**.** Cryo-electron tomograms of *ftsL** mutant. Time-lapse series were acquired with a rate of 7 fps in the compressed format MOV for visualization purposes. Green, cyan and magenta layers indicate segmented IM, PG and OM, respectively. Scale bars = 100 nm.

**Supplementary Video 6. Cell wall hydrolysis contributes to Z-ring condensation.** Three-dimensional maximum intensity projections rendered in Huygens (SVI) of indicated *E. coli* strain expressing Pal-mCherry (magenta) and ZipA-sfGFP (green) are shown. Video was rendered at 12 fps. Scale bar = 2µm.

**Supplementary Video 7: MreB filaments regularly pass either through or in direct proximity of the cell division site.** A three-minute SIM-TRIF time-lapse series of MreB-sw-mNeonGreen (green) was overlayed over a 3D-SIM image of Pal-mCherry (magenta) and bright field reference image. On the right-side tracking results from TrackMate are overlayed. Video was rendered at 12 fps. Scale bar = 1 µm. **Videos available upon request.**
